# Small-RNA Profiling Links 5’-tRNA Halves to Post-therapeutic Disease Persistence and Poor Patient Survival in Glioblastoma

**DOI:** 10.64898/2026.07.14.738483

**Authors:** Monima Anam, Taylor L Schanel, Sophia Dunlap, Mostafa Mohamed, Eun-Young Erin Ahn, Christopher D Willey, Zhangli Su

**Author notes:** These authors contributed equally (Taylor L Schanel, Sophia Dunlap, Eun-Young Erin Ahn, Christopher D Willey).

## Abstract

Glioblastoma (GBM) is a highly lethal brain cancer with limited therapeutic durability, where the majority of patients develop recurrent or persistent disease after standard chemoradiotherapy. Meanwhile, tRNA-derived fragments (tRFs) have become increasingly relevant to cancer biology; however, their clinical relevance in GBM remains undefined. Here, we report that a specific family of tRFs, 5’-tRNA halves (tiR5s) dominates the small RNA landscape of GBM patient tumors and associates with worse overall survival, post-therapeutic disease persistence, and pro-invasive proteogenomic pathways across two independent GBM patient cohorts. This association between elevated tiR5 levels and therapeutic resistance re-emerges in radiation-resistant GBM xenograft models. Our findings reveal that tiR5s are an underappreciated molecular feature of highly aggressive GBM tumors, supporting further investigation into their biological roles and prognostic utility in GBM.

**Highlights:**

- tiR5s are the predominant tRF family in primary GBM patient tumors
- Elevated tiR5 expression distinguishes primary GBM tumors that develop persistent disease after first-line therapy
- Radiation-resistant GBM PDX models show elevated tiR5 expression
- Elevated tiR5 expression associates with poor overall patient survival and pro-invasive molecular programs in GBM patient tumors

**Graphical Abstract:** 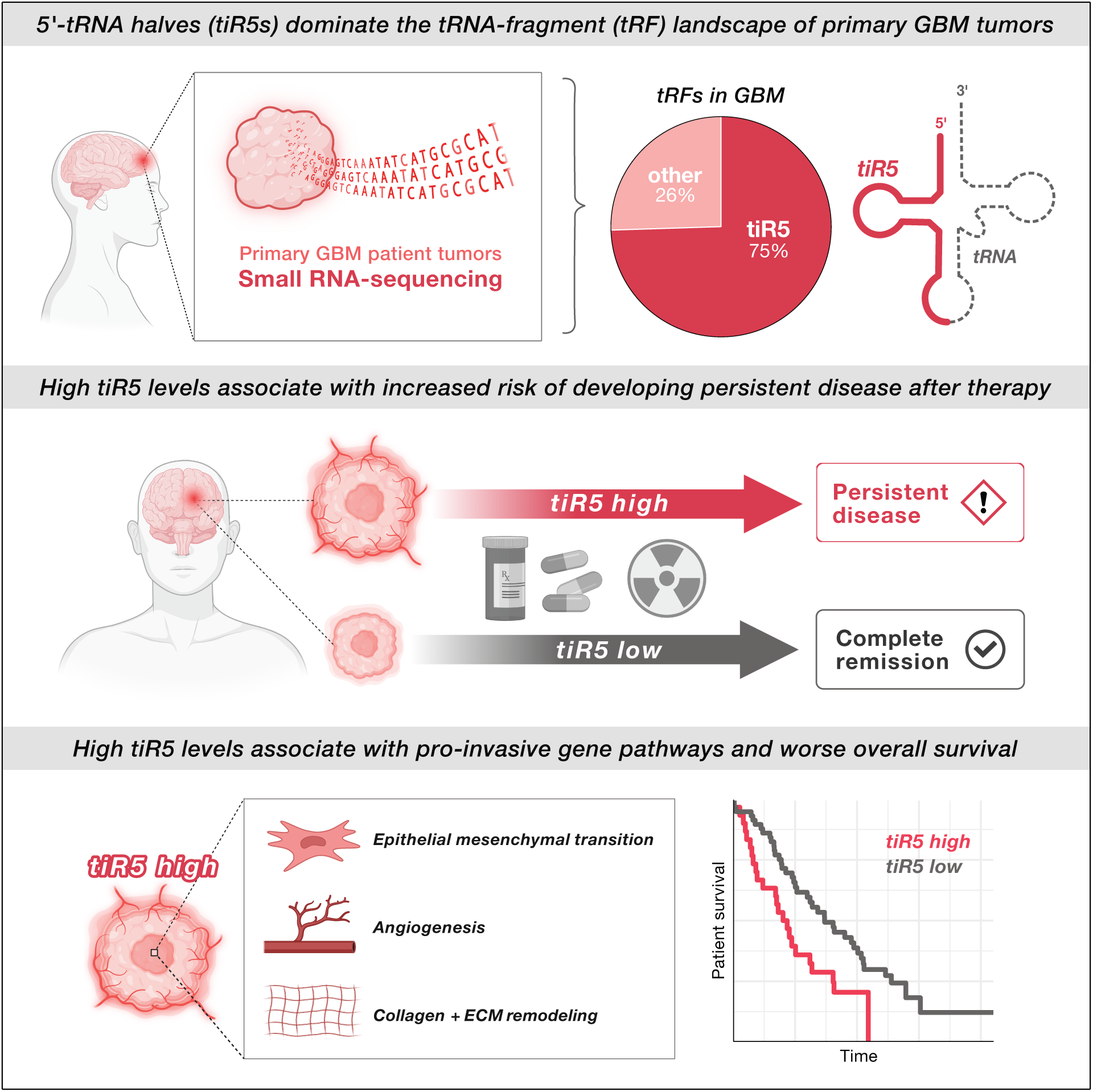

## Introduction

Glioblastoma (GBM) is a high-grade diffuse glioma, a brain tumor arising from glial or glial precursor cells, and one of the most common and lethal primary brain cancers in adults. Standard treatment consists of maximal safe surgical resection followed by radiotherapy with concurrent and adjuvant chemotherapy using temozolomide (TMZ). Despite this intensive treatment approach, median survival remains approximately 12-15 months after diagnosis, and recurrence is nearly inevitable.^1–4^ This therapeutic plateau is shaped in part by the profound heterogeneity of GBM, with variation across patients and within individual tumors at the genomic, histologic, and microenvironmental levels. Established clinical and molecular features such as age, molecular subtype, chromosomal alterations, and MGMT promoter methylation status provide important prognostic information, but incompletely capture the biological programs shaping treatment resistance and survival.^5–7^ Defining additional molecular features of GBM heterogeneity is therefore essential for improving tumor stratification and identifying biological pathways associated with aggressive disease.

Across cancers, small noncoding RNAs have been recognized as significant contributors to tumor heterogeneity. Among these, tRNA-derived fragments (tRFs), a class of small noncoding regulatory RNAs generated through enzymatic cleavage of tRNAs, have garnered attention due to their unexpected biological significance in cancer and disease.^8,9^ While tRNAs canonically deliver amino acids to the ribosome, tRFs have distinct regulatory functions unrelated to amino acid transfer.^10,11^ tRFs are classified by the region of the parental tRNA from which they are cleaved and the length of the resulting fragment. In general, tRFs act through diverse regulatory mechanisms, including Argonaute-RISC-associated repression, modulation of mRNA stability and translation, RNA-binding protein interactions, and cellular stress-adaptation.^11–18^ These tRF-dependent regulatory processes are directly applicable to cancer biology. Across multiple cancer types, tRFs have been shown to impact cell proliferation, migration, invasion, and metastatic progression in preclinical models.^19–23^ Additional preclinical studies have functionally linked tRFs to cancer therapeutic resistance.^24–27^ In parallel, clinical profiling has identified multiple tRF candidates that serve as prognostic biomarkers in cancer.^26,28–32^ Together, these studies establish tRFs as functional cancer-associated small RNAs, motivating further investigation of their clinical utility and biological relevance in GBM.

Among tRF classes, 5′ tRNA halves (tiR5s) are especially relevant to stress-adapted cancer states. More broadly, tRNA halves (tiRs) are ∼30-35 nucleotide fragments generated by cleavage of tRNAs near the anticodon loop. These tiRs were initially characterized as stress-responsive small RNAs induced by oxidative stress, nutrient deprivation, heat shock, and UV irradiation.^13,33–35^ tiR5s in particular have been shown to promote stress-adaptation by repressing translation, promoting stress granule assembly, and preventing apoptosis in cellular models.^12–14^ These stress-response pathways are highly relevant to cancer, where tumor cells must withstand metabolic stress, hostile microenvironments, and therapeutic toxicity. Meanwhile, tiR5s have been reported as highly abundant and dysregulated tRF species across multiple cancer types, where tiR5s have been linked to tumor progression and signaling and highlighted as candidate prognostic biomarkers.^19,36–45^ This convergence of stress responsiveness, high abundance, and cancer-associated dysregulation makes tiR5s a compelling tRF family for focused analysis in GBM.

Despite growing interest in tRFs across cancers, limited research has investigated tRFs and tiR5s in GBM. One study observed abundant tRFs in glioma stem cell-derived extracellular RNA, and two subsequent studies identified differentially expressed tRFs between GBM and low-grade glioma (LGG) patient tumors, suggesting that tRFs are detectable and biologically relevant in glioma. However, the relationship between tRF expression and clinically relevant heterogeneity within primary GBM, including treatment response, molecular phenotype, and survival outcomes, remains poorly defined.^46–49^

Here, we performed small RNA profiling of primary patient GBM tumors and non-tumor brain samples taken from the Clinical Proteomic Tumor Analysis Consortium (CPTAC-3) GBM project.^50^ We identify broad remodeling of the GBM tRF landscape and find that tiR5s are the predominant tRF family in primary GBM tumors across two independent patient cohorts. We further demonstrate that elevated tiR5 expression is associated with persistent disease after first-line chemoradiotherapy, increases after experimental induction of radiation resistance in GBM patient-derived xenograft (PDX) models, and is linked to worse overall patient survival after adjustment for established clinical and molecular variables, with findings replicated across cohorts. Finally, integration of transcriptomic and proteomic data shows that tiR5-high tumors are associated with mitochondrial translation, epithelial-to-mesenchymal transition, angiogenesis, and extracellular matrix remodeling programs. Together, these findings identify tiR5s as an underrecognized small RNA feature associated with aggressive GBM biology, poor treatment response, and adverse patient outcomes.

## Results

### Small RNA profiling reveals coordinated differential regulation of tRF families in primary patient GBM tumors

To characterize the small RNA landscape of primary patient GBM, we annotated publicly available small RNA-sequencing data from the CPTAC-3 GBM discovery cohort, including detailed annotation of tRFs. The dataset included resected pre-treatment primary GBM tumors (n = 92) and age-matched post-mortem non-tumor brain samples from GTEx (n = 8), which were processed alongside tumor samples using the CPTAC-3 workflow. Cohort characteristics are reported in **Table 1**.

**Table 1.** CPTAC-3 GBM cohort characteristics.

| Characteristic | Discovery cohort | Confirmatory cohort |
| --- | --- | --- |
| Patient samples |  |  |
| Total, N | 100 | 67 |
| Primary GBM tumor, n | 92 | 67 |
| Non-tumor brain, n | 8 | 0 |
| Patient demographics |  |  |
| Median age in years (range) | 59 (24-88) | 65 (29-84) |
| Median body mass index (range) | 25.7 (18.4-38.3) | 25.0 (19.0-34.0) |
| Female, n (%) | 45 (45.0%) | 21 (31.3%) |
| Male, n (%) | 55 (55.0%) | 46 (68.7%) |

Across all samples, miRNAs comprised the largest fraction of mapped small RNA reads, representing an average of 61.4% of mapped small RNAs. tRFs were the second most abundant annotated small RNA biotype, representing an average of 21.7% of mapped reads across GBM tumors and non-tumor brain samples (**Fig. 1A**). These results support that tRFs are a prominent component of the small RNA transcriptome in both primary GBM and non-tumor brain samples.

**Figure 1.**
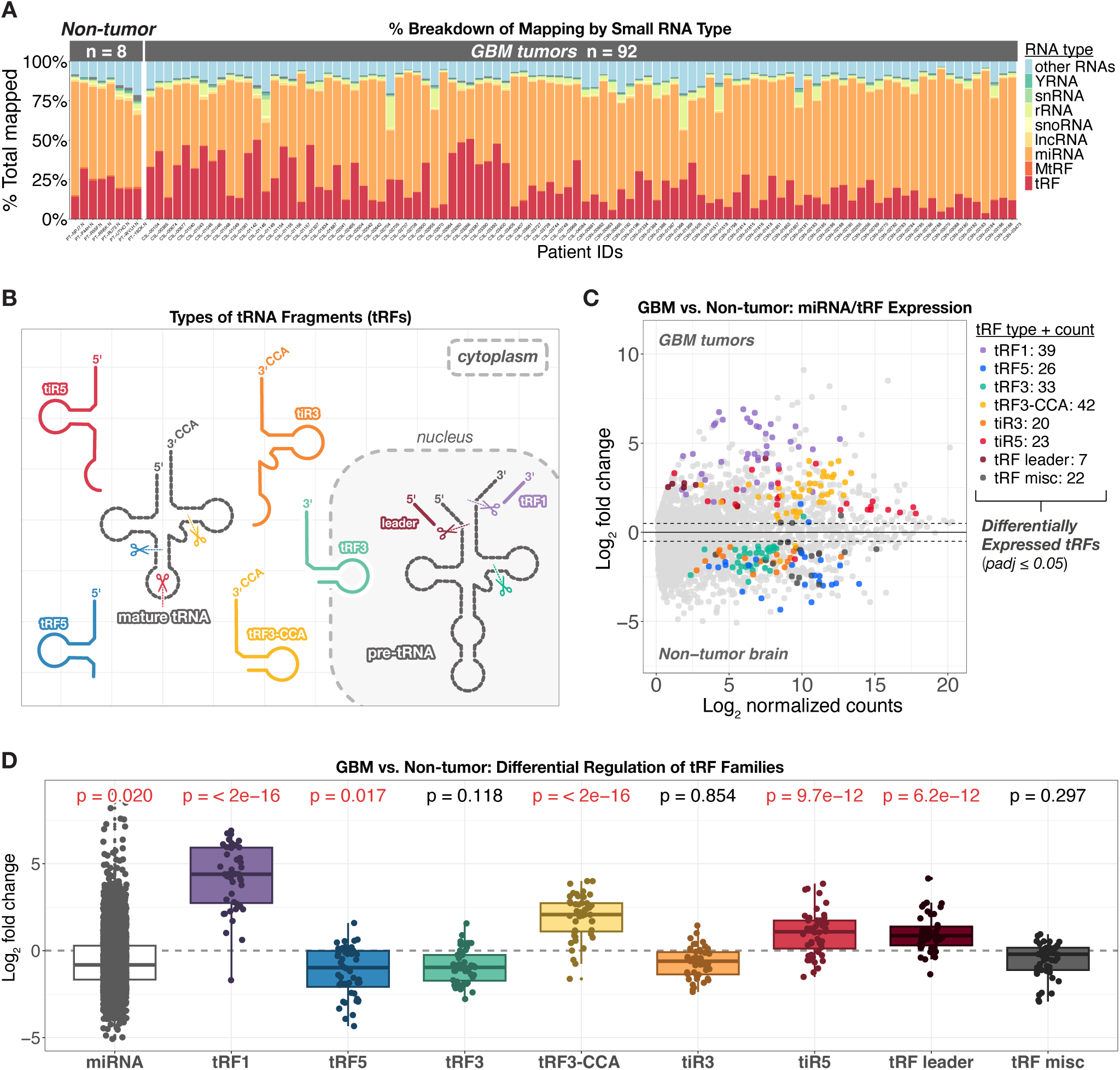
Small RNA profiling reveals differential regulation of tRF families in primary GBM tumors. **(A)** Stacked bar plot showing the percentage of mapped small RNA reads assigned to each RNA biotype in non-tumor brain samples (n = 8) and primary GBM tumors (n = 92). Each bar represents one patient sample, and each color represents one RNA biotype. **(B)** Schematic of major tRF families generated from mature tRNAs and pre-tRNAs. The schematic highlights the primary tRF families analyzed in this study; additional tRF subfamilies and cleavage variants are not shown. **(C)** MA plot showing differential expression of miRNAs and tRFs in GBM tumors compared with non-tumor brain samples. The x-axis shows mean DESeq2-normalized expression on a log_2_ scale, and the y-axis shows log_2_ fold change in GBM tumors versus non-tumor brain. Significantly differentially expressed tRFs are color-coded by tRF family using an adjusted p-value ≤ 0.05 and an absolute log_2_ fold change ≥ 0.5. Gray points indicate either tRFs that did not meet these thresholds, or miRNAs. **(D)** Boxplots showing log_2_ fold-change values for miRNAs and tRF families in GBM tumors compared with non-tumor brain samples. Each point represents one small RNA species. P-values above each boxplot indicate Wilcoxon tests comparing the fold-change distribution for that small RNA class with the fold-change distribution across all small RNAs shown in the panel. All p-values are adjusted for multiple hypothesis testing, and significant adjusted p-values ≤ 0.05 are highlighted in red.

We next performed differential expression analysis comparing GBM tumors with non-tumor brain samples. miRNAs showed widespread differential expression, with 436 miRNAs upregulated and 1,061 miRNAs downregulated in GBM tumors at an adjusted p-value ≤ 0.05 and absolute log_2_ fold change ≥ 0.5 (**Fig. S1**). We then focused on tRFs, which were grouped into eight distinct families according to fragment length and mapping pattern along parental tRNA sequences. Shorter fragments (< 30 nucleotides (nt)) included tRF5 and tRF3 species derived from the 5’ and 3’ ends of tRNAs, respectively, and tRF3-CCA species that retained the terminal 3’-CCA sequence found in mature tRNAs. Longer fragments (≥ 30 nt) arose from cleavage near the tRNA anticodon loop, producing 5’-and 3’-tRNA halves, referred to as tiR5s and tiR3s, respectively. Precursor tRNA (pre-tRNA) derived classes included tRF leader and tRF1 species cleaved from the pre-tRNA 5’-leader and 3’-trailer sequences, respectively. Fragments outside these categories were classified as tRF misc. (**Fig. 1B**).^11,51^ Individual tRF species are named according to the amino acid and anticodon of the parental tRNA, followed by the tRF family; for example, Gly-GCC-tiR5 denotes a tiR5 fragment cleaved from a parental glycine-carrying tRNA that contains a GCC anticodon.

Our analyses revealed that tRFs meeting differential expression criteria, defined as an adjusted p-value ≤ 0.05 and an absolute log_2_ fold change ≥ 0.5, showed coordinated family-level changes in GBM tumors compared with non-tumor brain samples. Differentially expressed tRFs exhibited coordinated, family-specific patterns of regulation. tRF1, tRF3-CCA, tiR5, and tRF leader species were predominantly upregulated in GBM tumors, whereas tRF5, tRF3, tiR3, and tRF misc species were predominantly downregulated (**Fig. 1C**; **Fig. S1**). This directional pattern was most pronounced for tRF1 and tRF leader species, with all 39 differentially expressed tRF1 species and all 7 differentially expressed tRF leader species increased in GBM tumors. tRF3-CCA and tiR5 species showed a similar pattern, with 40 of 42 differentially expressed tRF3-CCA species and 22 of 23 differentially expressed tiR5 species increased in GBM tumors. By contrast, most differentially expressed tRF5, tRF3, tiR3, and tRF misc species were decreased in GBM tumors compared to non-tumor brain samples (**Fig. 1C**; **Fig. S1**).

We next asked whether these directional patterns were limited to differentially expressed tRFs or reflected broader shifts across all annotated members of each tRF family. To address this, we compared the log_2_ fold-change distributions for all detected species within each tRF family, as well as all detected miRNAs, regardless of whether individual species met differential expression criteria (**Fig. 1D**). This analysis showed significant family-wide positive shifts for tRF1, tRF3-CCA, tiR5, and tRF leader species, consistent with increased abundance in GBM tumors. In contrast, tRF5 species showed a significant family-wide negative shift, consistent with reduced abundance in GBM tumors. miRNAs also showed an overall negative shift, consistent with previously reported GBM literature.^52^ These family-level shifts remained significant after multiple-testing correction at adjusted p-value ≤ 0.05, whereas tRF3, tiR3, and tRF misc families did not display significant family-wide shifts (**Fig. 1D**).

Together, these analyses identify broad remodeling of the small RNA transcriptome in primary GBM tumors. Beyond widespread miRNA dysregulation, tRF changes occurred in a coordinated family-specific pattern, with selective enrichment of tRF1, tRF3-CCA, tiR5, and tRF leader species and depletion of tRF5 species in GBM tumors. The full DESeq2 outputs for GBM tumor versus non-tumor comparisons are provided in **Dataset S1**. These findings suggest that tRF dysregulation in GBM is organized at the fragment-family level rather than distributed uniformly across tRF classes.

### tiR5s are the predominant tRF family in primary patient GBM tumors

Having observed the coordinated differential expression of tRF families in GBM tumors, we next asked whether a particular tRF family predominated in GBM tumor tissue. For these analyses, we used the CPTAC-3 GBM discovery cohort from the preceding tumor versus non-tumor analyses, together with the independent CPTAC-3 GBM confirmatory cohort (**Table 1**). The confirmatory cohort lacked matched non-tumor brain samples, precluding tumor versus non-tumor comparisons, but included primary GBM tumor small RNA profiles for independent evaluation of tumor-level tRF abundance. In the discovery cohort (N = 92), tiR5 species represented the dominant tRF class, accounting for an average of 74.5% of all mapped tRF reads. The remaining most abundant tRF families represented much smaller fractions, including tRF misc (14.3%), tRF3-CCA (4.6%), and tRF5 species (4.3%) (**Fig. 2A**). This tiR5 predominance was reproduced in the confirmatory cohort (N = 67), where tiR5 species again accounted for 74.5% of total tRF reads in GBM tumors (**Fig. S2A**).

**Figure 2.**
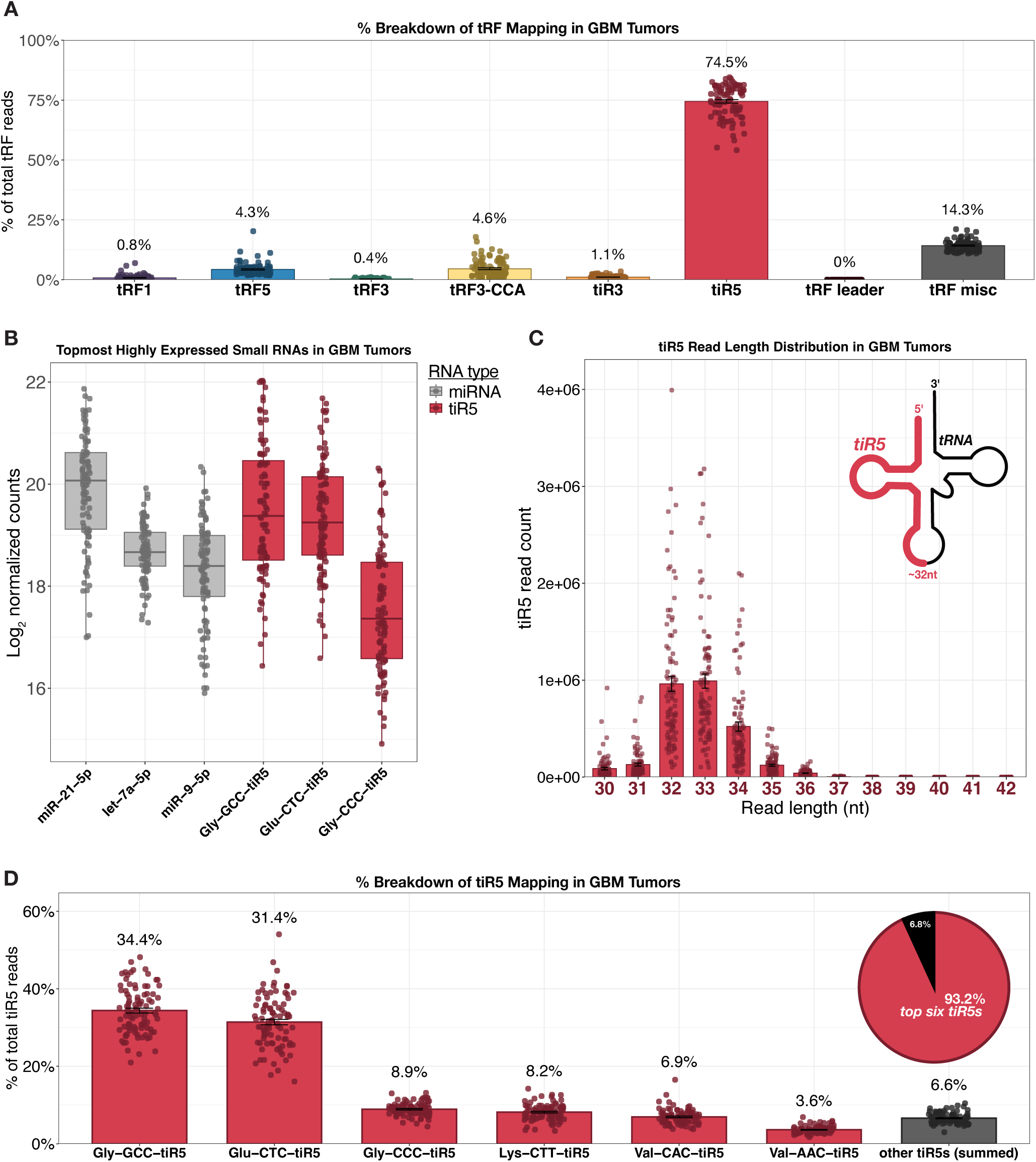
tiR5s are the predominant tRF family in primary GBM tumors. **(A)** Bar plot showing the percentage of mapped tRF reads assigned to each tRF family in primary GBM tumors (N = 92). Percentages above each bar indicate the mean percentage of total tRF reads assigned to each respective tRF family. Error bars represent SEM. **(B)** Boxplots showing the six most highly expressed small RNAs in GBM tumors. The y-axis shows the log_2_-transformed DESeq2-normalized counts for each small RNA. Three of these six most highly expressed small RNAs consist of tiR5 species. **(C)** Read-length distribution of tiR5-mapped reads in GBM tumors. The x-axis shows tiR5 read length in nucleotides, and the y-axis shows the total number of tiR5 reads at each length. Error bars represent SEM. The accompanying schematic illustrates a representative parental tRNA and the corresponding tiR5 fragment, highlighted in red. **(D)** Bar plot showing the percentage of total tiR5 reads assigned to individual tiR5 species in GBM tumors. Percentages above each bar indicate the mean percentage of total tiR5 reads assigned to each respective tiR5 species. Error bars represent SEM. The inset pie chart reports that the six most abundant individual tiR5 species account for 93.2% of total tiR5 reads, with all remaining tiR5 species grouped as “other tiR5s.”

We next compared tiR5 abundance with the most highly expressed small RNAs detected in GBM tumors using the discovery cohort. Among the six most abundant small RNAs, three were well-established highly expressed miRNAs: miR-21-5p, let-7a-5p, and miR-9-5p, and three were tiR5 species: Gly-GCC-tiR5, Glu-CTC-tiR5, and Gly-CCC-tiR5 (**Fig. 2B**). These tiR5-mapped reads clustered at approximately 32-33 nucleotides, consistent with the length of 5’-tRNA halves generated by tRNA cleavage near the anticodon loop (**Fig. 2C**). Overall, these findings identify tiR5s as the most abundant tRF family in primary GBM tumors.

We detected 48 unique tiR5 species in GBM tumors; however, tiR5 abundance was concentrated among the top six most highly expressed tiR5s: Gly-GCC-tiR5, Glu-CTC-tiR5, Gly-CCC-tiR5, Lys-CTT-tiR5, Val-CAC-tiR5, and Val-AAC-tiR5. Together, these six species accounted for 93.2% of total tiR5 reads in the discovery cohort, whereas all remaining tiR5 species accounted for 6.8% of tiR5 reads (**Fig. 2D**). Likewise, the same six tiR5 species accounted for 94.5% of total tiR5 reads in the confirmatory cohort (**Fig. S2B**). These six dominant tiR5 species were strongly co-expressed across tumors in both cohorts. In the discovery cohort, pairwise Pearson correlation coefficients ranged from 0.90 to 0.98, with adjusted p-values < 0.001 for all comparisons (**Fig. S2C**). In the confirmatory cohort, pairwise Pearson correlation coefficients ranged from 0.85 to 0.97, again with adjusted p-values < 0.001 for all comparisons (**Fig. S2D**). At the tRF family level, total tiR5 expression was positively correlated with tRF5, tiR3, and tRF misc abundance in both cohorts (adjusted p < 0.001; **Fig. S2E-F**). Total tiR5 expression also showed an inverse correlation with tRF3-CCA abundance in both cohorts (adjusted p < 0.01; **Fig. S2E**). Overall, these results show that tiR5 abundance is concentrated within a small, highly co-expressed set of dominant tiR5 species across two independent GBM cohorts.

Finally, we assessed whether variation in total tiR5 expression could be explained by differences in tumor tissue composition. Total tiR5 expression was not significantly correlated with stromal score (R = 0.038, p = 0.72), immune score (R = 0.027, p = 0.80), or tumor purity (R = −0.046, p = 0.67) (**Fig. S2G**). These results argue against stromal content, immune content, or tumor purity as major drivers of intertumoral variation in tiR5 expression. Together, these analyses identify tiR5s as the predominant tRF family in primary GBM tumors, driven by a set of small, highly co-expressed tiR5 species that are reproduced across independent patient cohorts.

### Elevated tiR5 expression is associated with persistent disease after first-line therapy in primary patient GBM tumors

We next asked whether tumor tRF expression was associated with clinical and molecular features of GBM. For these analyses, we used small RNA profiles and available clinical annotations from the CPTAC-3 GBM discovery cohort together with the independent CPTAC-3 GBM confirmatory cohort. In both cohorts, primary tumors were collected at surgical resection before first-line therapy and profiled by small RNA-sequencing. Patients then received standard first-line chemoradiotherapy, after which treatment response was classified as persistent disease or complete remission (**Fig. 3A**).

**Figure 3.**
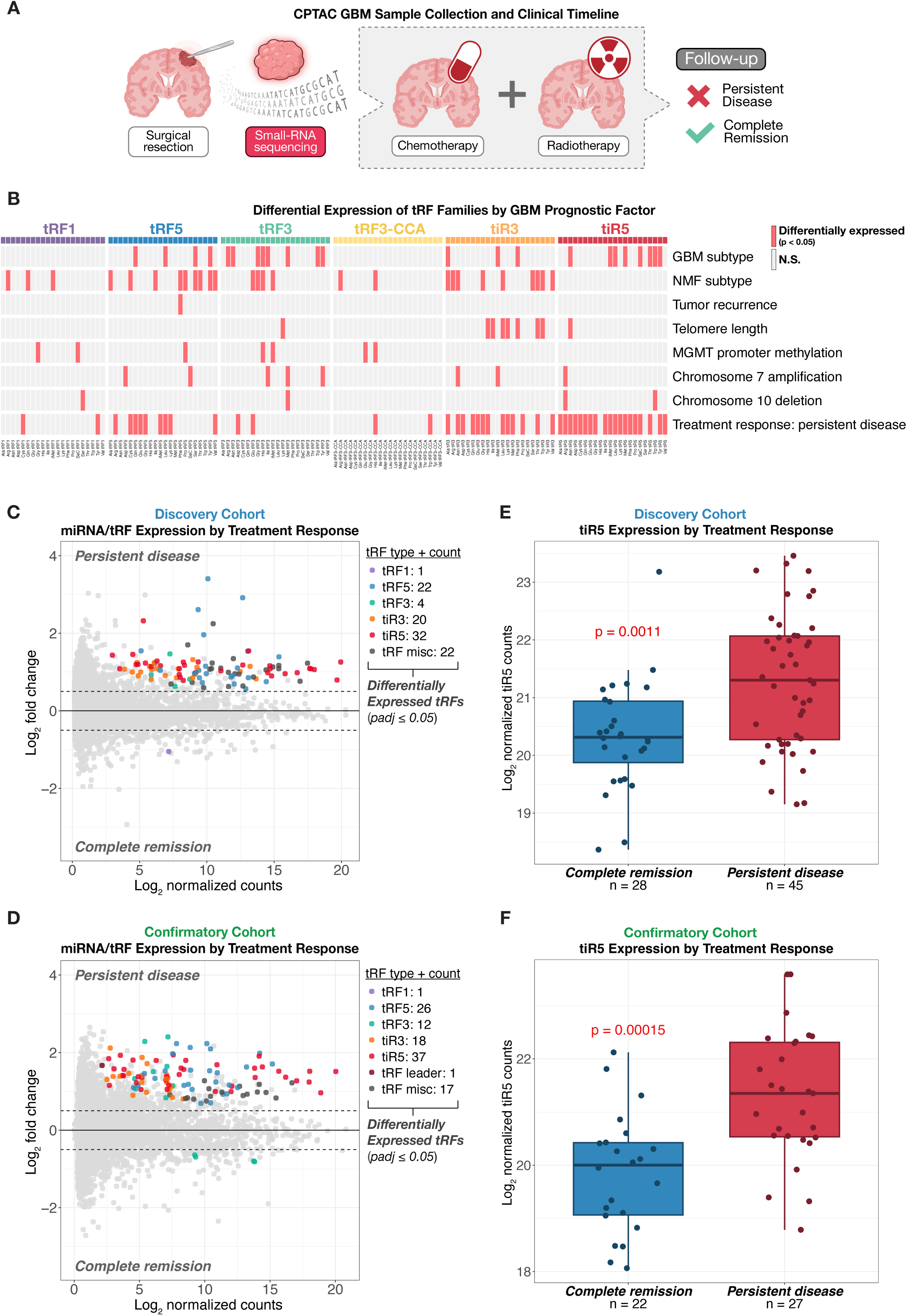
Elevated tiR5 expression is associated with persistent disease after first-line GBM therapy. **(A)** Schematic showing CPTAC GBM sample collection and clinical treatment timeline. Primary GBM tumors were collected at surgical resection and profiled by small RNA-sequencing pre-treatment. Treatment response was assessed after first-line chemoradiotherapy, where tumors were then classified as showing persistent disease or complete remission. **(B)** Heatmap showing differential expression of tRF species across GBM clinical and molecular features. Columns represent individual tRF species grouped by tRF family, and rows represent GBM-relevant clinical or molecular features. Red cells indicate tRF species that are significantly differentially expressed for a given feature using an adjusted p-value ≤ 0.05. Gray cells indicate non-significant comparisons. **(C-D)** MA plots showing differential expression of miRNAs and tRFs in GBM tumors that went on to develop persistent disease versus tumors that went into complete remission following first-line therapy. The x-axis shows mean DESeq2-normalized expression on a log_2_ scale, and the y-axis shows log_2_ fold change for persistent disease versus complete remission comparisons. Significantly differentially expressed tRFs are highlighted and color-coded by tRF family using an adjusted p-value ≤ 0.05 and an absolute log_2_ fold change ≥ 0.5, indicated by the dashed lines. Gray points indicate either tRFs that did not meet these thresholds, or miRNAs. Discovery **(C)** and confirmatory **(D)** patient cohorts are shown separately. **(E-F)** Boxplots showing total tiR5 expression in complete remission versus persistent disease tumor groups. The y-axis shows log_2_-transformed DESeq2-normalized tiR5 counts, where each point represents one patient sample. P-values represent Wilcoxon tests comparing tiR5 expression between complete remission and persistent disease tumor groups. Discovery **(E)** and confirmatory **(F)** patient cohorts are shown separately.

We first screened tRF species in the discovery cohort for associations with GBM-relevant clinical and molecular variables, including GBM subtype, non-negative matrix factorization (NMF)-defined molecular subtype, tumor recurrence, telomere length, MGMT promoter methylation, chromosome 7 amplification, chromosome 10 deletion, and treatment response. This initial analysis was limited to the discovery cohort due to incomplete annotation of these features in the confirmatory cohort. Differentially expressed tRF species were observed across multiple clinical and molecular features (**Fig. 3B**). Treatment response was then prioritized for subsequent analysis, as nearly all tiR5 species were increased in tumors that went on to develop persistent disease compared with tumors that underwent complete remission following first-line therapy.

We then examined treatment response associations by tRF family across both cohorts. A total of 48 unique tiR5 species were detected in both cohorts (**Fig. 3C-D**). In the discovery cohort (N = 73), 32 of these tiR5 species were significantly upregulated in tumors that went on to develop persistent disease, making tiR5s the largest differentially expressed tRF group by species count (**Fig. 3C**). This pattern was reproduced in the confirmatory cohort (N = 49), where 37 of 48 tiR5 species were significantly upregulated in tumors that went on to develop persistent disease, again representing the largest treatment-associated tRF group (**Fig. 3D**). No tiR5 species were significantly downregulated in tumors that developed persistent disease in either cohort. Although other tRF families, including tRF5, tiR3, and tRF misc, also showed differential expression by treatment response, tiR5s showed the most consistent treatment-associated upregulation across both cohorts and represented the most abundant tRF species.

We next compared total tiR5 expression between treatment-response groups. Total tiR5 abundance was significantly higher in tumors that went on to develop persistent disease than in tumors that underwent complete remission in both the discovery cohort (fold change = 1.94, p = 0.0011) and the confirmatory cohort (fold change = 2.71, p = 0.00015) (**Fig. 3E-F**). In the discovery cohort, where broader clinical and molecular annotations were available, a patient-level heatmap of total tiR5 expression and the six most abundant tiR5 species showed that treatment response aligned more clearly with tiR5-high tumors than did the other annotated features (**Fig. S3A**). Together, these results identify elevated tiR5 expression as a reproducible feature of GBM tumors that go on to develop persistent disease after first-line therapy. The full DESeq2 outputs for persistent disease versus complete remission comparisons are provided in **Dataset S2**.

### tiR5 expression improves prediction of persistent disease after first-line therapy in primary patient GBM tumors

Having observed higher tiR5 expression in GBM tumors that went on to develop persistent disease, we next asked whether tiR5 expression improved classification of treatment response after first-line therapy. We performed repeated cross-validated ROC analysis comparing clinical models alone, clinical models plus total tiR5 expression, and total tiR5 expression alone. Model performance was summarized by the area under the ROC curve (AUC), where higher AUC values indicate better separation between tumors that developed persistent disease and tumors that underwent complete remission.

Two baseline clinical models were used for ROC analysis. Clinical model A included age, sex, BMI, and MGMT promoter methylation status from clinical metadata. The confirmatory cohort lacked this clinical metadata, so model A was applied solely to the discovery cohort (**Fig. 4B**). To compare the discovery and confirmatory cohorts using the same variables, we created clinical model B, which used age, sex, BMI, and array-based MGMT promoter methylation status as a surrogate MGMT variable (**Fig. 4A**). Clinical model B was then evaluated in both cohorts (**Fig. 4C-D**).

**Figure 4.**
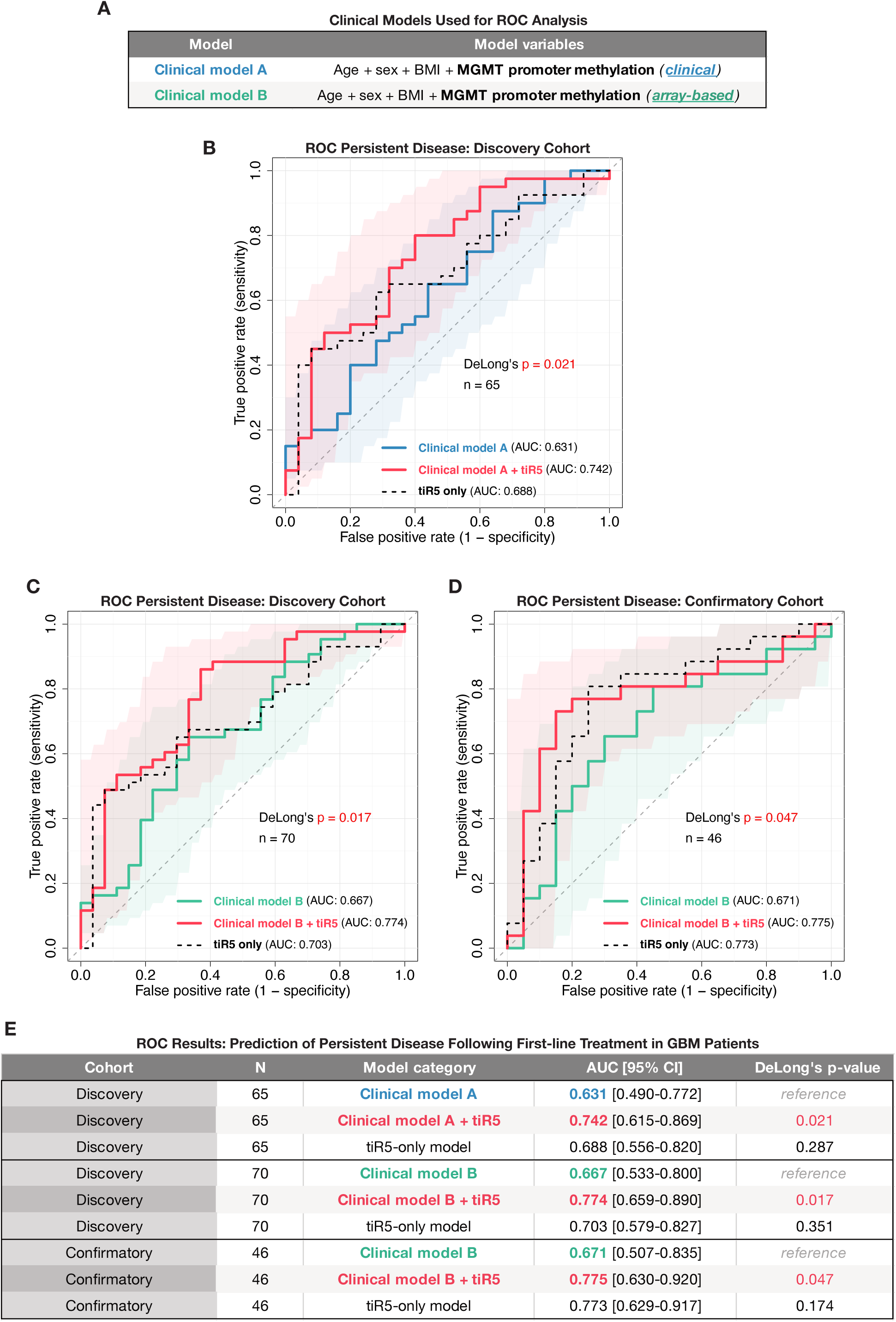
tiR5 expression improves prediction of persistent disease after first-line GBM therapy. **(A)** Summary of clinical models used for ROC analysis. Clinical model A includes age, sex, BMI, and clinically annotated MGMT promoter methylation status. Clinical model B includes age, sex, BMI, and array-based MGMT promoter methylation status. Clinical model A was evaluated in the discovery cohort, where clinical MGMT status information was available. Clinical model B was evaluated in both the discovery and confirmatory cohorts to enable cross-cohort comparison using a consistent array-derived MGMT classification. **(B)** Repeated cross-validated ROC analysis comparing clinical model A, clinical model A plus tiR5 expression, and tiR5 expression alone for prediction of persistent disease following first-line therapy in the discovery cohort. Shaded regions indicate 95% confidence intervals across cross-validation repeats. DeLong’s p-value reflects one-sided comparison between clinical model A and clinical model A plus tiR5 expression. **(C-D)** Repeated cross-validated ROC analysis comparing clinical model B, clinical model B plus tiR5 expression, and tiR5 expression alone for prediction of persistent disease following first-line therapy. Shaded regions indicate 95% confidence intervals across cross-validation repeats. DeLong’s p-value reflects one-sided comparison between clinical model B and clinical model B plus tiR5 expression. Discovery **(C)** and confirmatory **(D)** cohorts are shown separately. **(E)** Summary table reporting cohort, sample size, model category, AUC with 95% confidence interval, and DeLong’s p-value (one-sided). Clinical model A or B serves as the reference model for each comparison. Significant DeLong’s p-values ≤ 0.05 are highlighted in red.

In the discovery cohort (N = 65), total tiR5 expression improved prediction of persistent disease beyond baseline clinical variables. For clinical model A, adding tiR5 expression increased the AUC from 0.631 (95% CI, 0.490-0.772) to 0.742 (95% CI, 0.615-0.869), representing a significant improvement in discriminative performance by DeLong’s test (one-sided p = 0.021). Total tiR5 expression alone achieved an AUC of 0.688 (95% CI, 0.556-0.820), which was not significantly different from clinical model A (one-sided p = 0.287), indicating that tiR5 expression alone performed comparably to the baseline clinical model (**Fig. 4B** and **E**).

This improvement was reproduced using clinical model B in the discovery cohort (N = 70), where adding tiR5 expression increased the AUC from 0.667 (95% CI, 0.533-0.800) to 0.774 (95% CI, 0.659-0.890; p = 0.017). Total tiR5 expression alone achieved an AUC of 0.703 (95% CI, 0.579-0.827), again performing comparably to the baseline clinical model (one-sided p = 0.351; **Fig. 4C** and **E**). In the independent confirmatory cohort (N = 46), clinical model B showed the same pattern; adding tiR5 expression increased the AUC from 0.671 (95% CI, 0.507-0.835) to 0.775 (95% CI, 0.630-0.920), representing a significant improvement in discriminative performance by DeLong’s test (one-sided p = 0.047). Total tiR5 expression alone achieved a similar AUC of 0.773 (95% CI, 0.629-0.917) and was not significantly different from clinical model B (one-sided p = 0.174; **Fig. 4D-E**). Overall, these results demonstrate that total tiR5 expression improves prediction of persistent disease after first-line therapy beyond baseline clinical models across two independent GBM cohorts. The improved performance of tiR5-augmented models supports total tiR5 expression as a candidate biomarker for identifying primary GBM tumors at higher risk of post-treatment disease persistence.

### tiR5 expression increases following induction of radiation resistance in GBM PDX models

Given the association between elevated tiR5 expression and persistent disease following first-line GBM therapy, we next asked whether tiR5 expression was altered in an experimental model of acquired therapeutic resistance. To address this, we performed small RNA-sequencing of seven GBM PDX models in which baseline tumors were subjected to repeated radiation exposure to generate matched radiation-resistant tumors (**Fig. 5A**).^53^ This paired design enabled comparison of tiR5 expression before and after induction of radiation resistance within the same PDX models.

**Figure 5.**
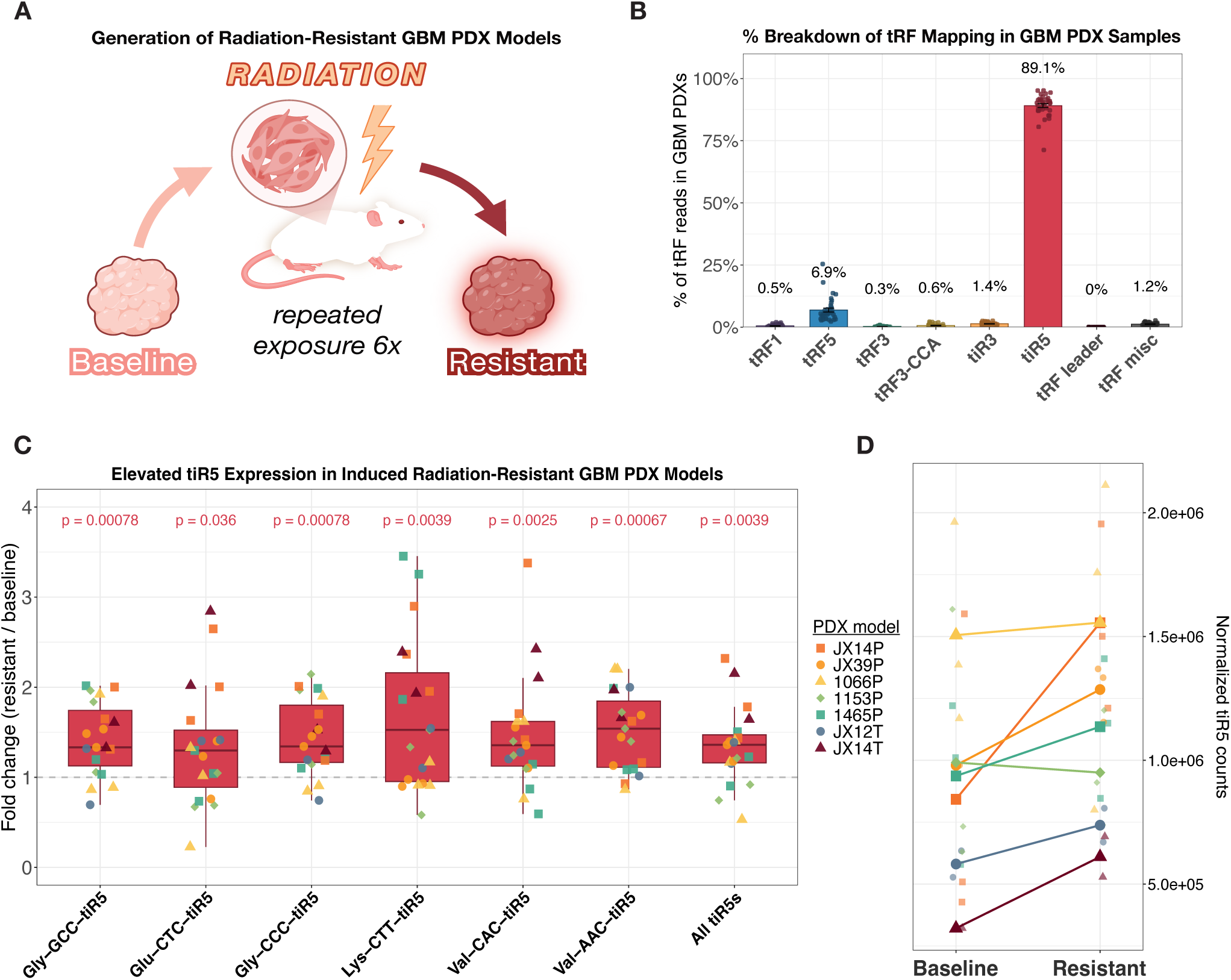
tiR5 expression increases following induction of radiation resistance in GBM PDX models. **(A)** Schematic showing the generation of radiation-resistant GBM PDX models. Baseline GBM PDX tumors were serially exposed to radiation to enrich for radiation-resistant cells. Baseline and resistant PDX samples were then profiled using small RNA-sequencing. **(B)** Bar plot showing the percentage of mapped tRF reads assigned to each tRF family in GBM PDX models. Each point represents one PDX sample (N = 35). Percentages above each bar indicate the mean percentage of total tRF reads assigned to each tRF family. **(C)** Boxplots showing resistant-to-baseline fold change for the six most abundant individual tiR5 species alongside total tiR5 expression in matched GBM PDX models. Each point represents one matched PDX model pair. P-values above each boxplot indicate Wilcoxon tests comparing each fold-change distribution against a null fold change of 1. All p-values are adjusted for multiple hypothesis testing, and significant adjusted p-values ≤ 0.05 are highlighted in red. **(D)** Line plot showing total tiR5 expression for each PDX model at baseline and after induction of radiation resistance. For each model, small points represent individual PDX samples or replicates. Larger points represent the mean total tiR5 expression across PDX replicates within each condition.

Consistent with the primary GBM tumor results from CPTAC-3 patients, tiR5s represented the predominant tRF family in GBM PDX samples. Across PDX tumors, tiR5 species accounted for 89.1% of mapped tRF reads on average, whereas tRF5, tiR3, and tRF misc species accounted for 6.9%, 1.4%, and 1.2% of mapped tRF reads, respectively. Other tRF families each represented less than 1% of mapped tRF reads (**Fig. 5B**). These results show that tiR5 predominance is conserved in GBM PDX models.

We next compared tiR5 expression between baseline and radiation-resistant PDX tumors by calculating resistant-to-baseline fold change for each matched PDX model. For each tiR5 species, fold-change values across the seven matched PDX pairs were tested against a null fold change of 1, corresponding to no change between resistant and baseline tumors. Total tiR5 expression increased after induction of radiation resistance, with a mean resistant-to-baseline fold change of 1.34 across PDX pairs (adjusted p = 0.0039; **Fig. 5C-D**). The six most abundant tiR5 species identified in primary GBM tumors also showed significant increases, with mean fold changes across PDX pairs ranging from 1.34 to 1.68 (adjusted p = 0.036 to 0.00067; **Fig. 5C**). Although the magnitude of induction varied across individual PDX models, most matched models showed higher total tiR5 expression after induction of radiation resistance (**Fig. 5D**).

Together, these findings show that tiR5s are the predominant tRF family in GBM PDX models and that total tiR5 expression, as well as individual expression of the top six most abundant tiR5s, significantly increases after experimental induction of radiation resistance. These results provide orthogonal support for the association between elevated tiR5 expression and poor response to first-line therapy observed in primary patient GBM tumors.

### tiR5-high primary patient GBM tumors exhibit proteomic enrichment of mitochondrial translation pathways

Having established that elevated tiR5 expression is associated with treatment resistance in both patient tumors and radiation-resistant PDX models, we next sought to identify the molecular pathways associated with tiR5-high GBM tumors. For proteome-wide pathway analysis, we used MS1-based log_2_ protein intensity values from the CPTAC-3 GBM discovery cohort (N = 92),^54^ which provided protein abundance measurements suitable for pathway-level enrichment analysis. Tumors were stratified into tiR5-high and tiR5-low groups based on mean total tiR5 expression, and proteins were ranked by differential abundance between groups for Gene Set Enrichment Analysis (GSEA) using Reactome terms. This analysis was restricted to the discovery cohort due to the lack of MS1-based protein abundance values in the confirmatory cohort.

The four highest-ranked Reactome pathways enriched in tiR5-high tumors were all related to mitochondrial translation: mitochondrial translation elongation (NES = 2.86, adjusted p = 1.5 x 10^-8^), mitochondrial translation initiation (NES = 2.85, adjusted p = 1.5 x 10^-8^), mitochondrial translation (NES = 2.84, adjusted p = 1.5 x 10^-8^), and mitochondrial translation termination (NES = 2.84, adjusted p = 1.5 x 10^-8^) (**Fig. 6A**). Other top enriched pathways included RNA processing, mRNA splicing, extracellular matrix organization, and translation, but mitochondrial translation represented the strongest pathway-level signal in tiR5-high tumors.

**Figure 6.**
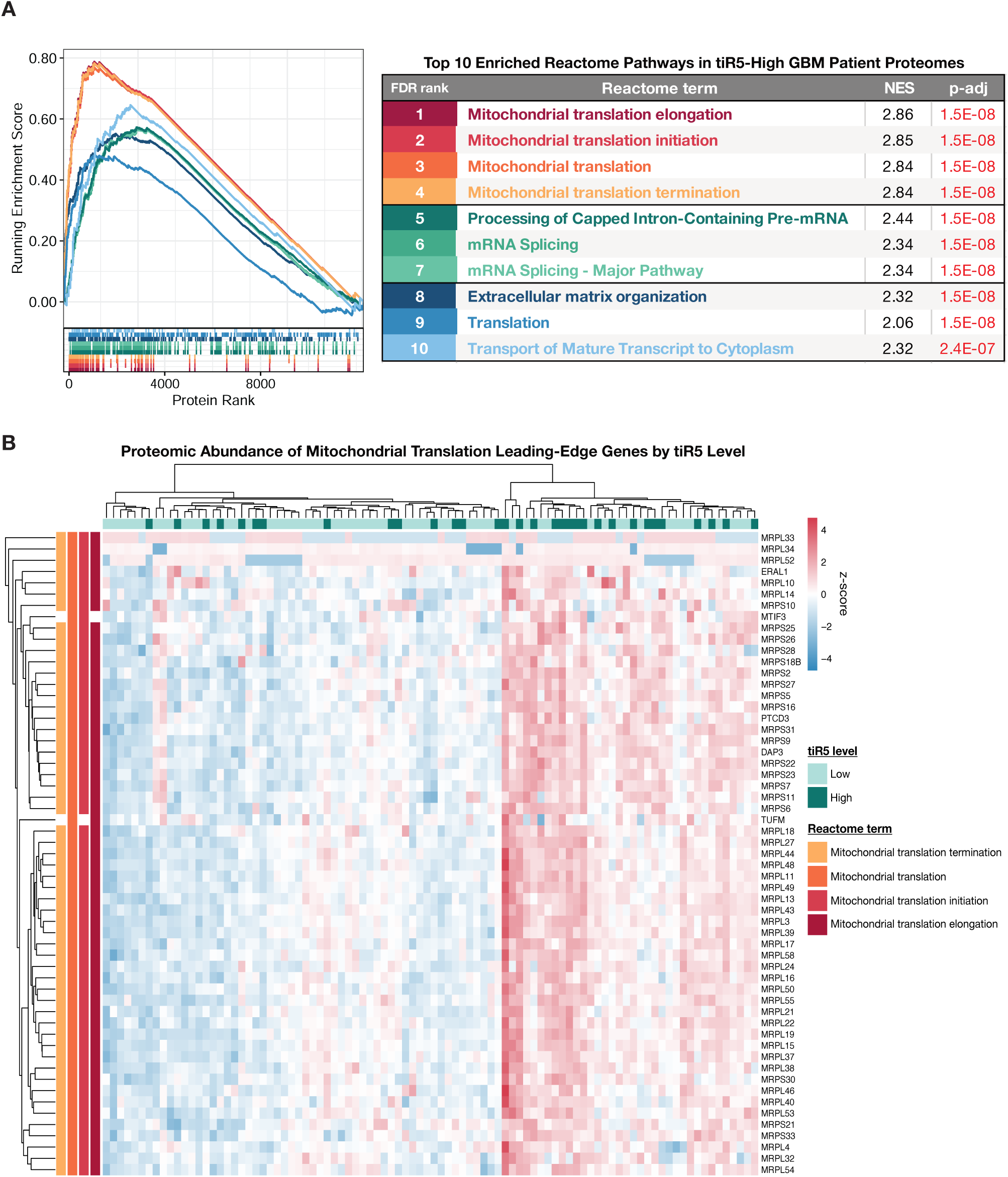
tiR5-high GBM patient tumors exhibit proteomic enrichment of mitochondrial translation pathways. **(A)** GSEA enrichment curves showing the top 10 Reactome pathways enriched in tiR5-high GBM tumors using patient tumor proteomics data. tiR5-high (n = 32) and tiR5-low (n = 60) groups were defined by mean tiR5 expression. Pathways are ranked by adjusted p-value and color-coded consistently between the enrichment curves, barcode tracks, and results table. The accompanying table reports the Reactome term, normalized enrichment score (NES), and adjusted p-value. Positive NES values indicate enrichment in tiR5-high tumors. **(B)** Heatmap showing proteomic abundance of mitochondrial translation leading-edge proteins from the enriched Reactome pathways in panel A. Columns represent each individual patient-tumor (N = 92), and rows represent leading-edge proteins. Samples are annotated by tiR5 expression group. Protein abundance is shown as a row-scaled z-score across samples.

We then visualized the leading-edge proteins contributing to the mitochondrial translation enrichment signal as a patient-level heatmap. Samples with higher abundance of mitochondrial ribosomal proteins and related mitochondrial translation factors included many tiR5-high tumors, consistent with the positive GSEA enrichment observed in tiR5-high tumors (**Fig. 6B**). These results identify mitochondrial translation machinery as a prominent proteomic feature of tiR5-high primary GBM tumors.

### tiR5-high patient GBM tumors show concordant transcriptomic and proteomic enrichment of epithelial mesenchymal transition (EMT), angiogenesis, and extracellular matrix (ECM) remodeling pathways

We next asked which tiR5-high-associated molecular pathways were conserved across bulk RNA and protein expression layers. Using the CPTAC-3 GBM discovery cohort (N = 92), tumors were stratified into tiR5-high and tiR5-low groups based on mean total tiR5 expression. Parallel GSEA was then performed using RNA-seq and MS1-based proteomic data and significantly enriched (adjusted p ≤ 0.05) MSigDB Hallmark and Reactome pathways were compared across both omics layers to identify concordant tiR5-high-associated pathways. This analysis was again restricted to the discovery cohort due to the lack of MS1-based protein abundance values in the confirmatory cohort.

Concordant positively enriched pathways centered on EMT, angiogenesis, and ECM remodeling (**Fig. 7A**). Among Hallmark pathways, two pathways were significantly positively enriched in tiR5-high tumors across both transcriptomic and proteomic layers: EMT (transcriptomic NES = 1.39, adjusted p = 1.4 x 10^-2^; proteomic NES = 1.91, adjusted p = 3.5 x 10^-5^) and angiogenesis (transcriptomic NES = 2.03, adjusted p = 9.4 x 10^-4^; proteomic NES = 1.70, adjusted p = 4.2 x 10^-2^) (**Fig. 7B**). Reactome GSEA identified 12 concordantly positively enriched pathways, with shared enrichment dominated by terms surrounding ECM organization, collagen biology, and cell-matrix interactions (**Fig. 7C**). Although these pathways overlap with mesenchymal-like GBM biology, total tiR5 expression did not significantly vary across classical, proneural, and mesenchymal tumors (Kruskal-Wallis p = 0.075; **Fig. S3B**), suggesting that the tiR5-high-associated pro-invasive signatures are not simply explained by enrichment of mesenchymal-subtype GBM. Together, these concordant Hallmark and Reactome results indicate that tiR5-high tumors are associated with pro-invasive and angiogenic programs across RNA and protein expression layers.

**Figure 7.**
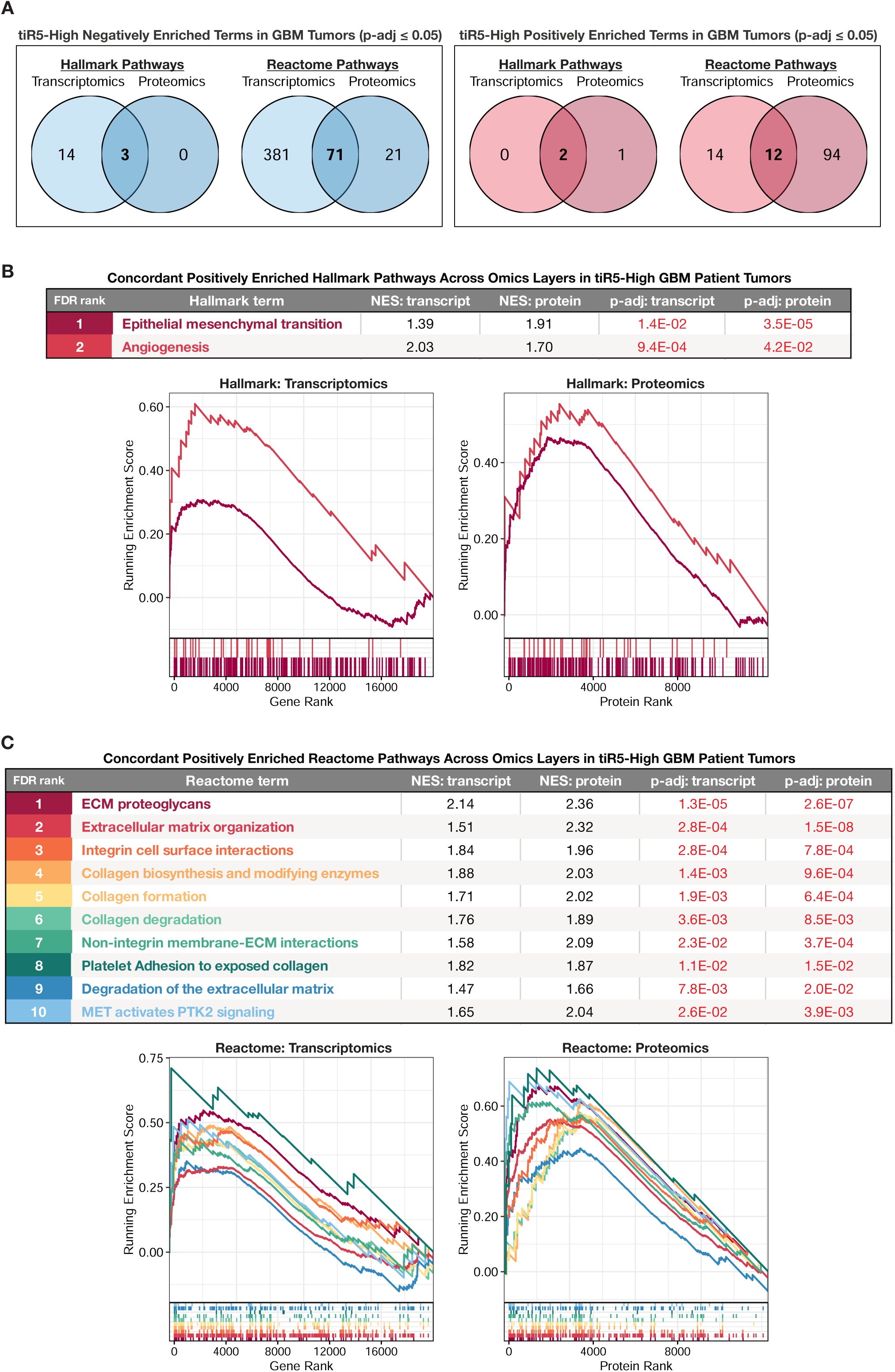
tiR5-high GBM patient tumors exhibit concordant transcriptomic and proteomic enrichment of EMT, angiogenesis, and ECM remodeling pathways. **(A)** Venn diagrams showing overlap of significantly enriched pathways (p-adj ≤ 0.05) between transcriptomic and proteomic GSEA analysis in tiR5-high GBM tumors. Negatively enriched pathways are shown on the left, and positively enriched pathways are shown on the right. MSigDB Hallmark and Reactome pathway collections are shown separately. **(B)** Concordant positively enriched MSigDB Hallmark pathways in tiR5-high GBM tumors across transcriptomic and proteomic datasets. The table reports the Hallmark term name, normalized enrichment score (NES), and adjusted p-value for each pathway. Pathways are color-coded consistently between the table, enrichment curves, and barcode tracks. **(C)** Concordant positively enriched Reactome pathways in tiR5-high GBM tumors across transcriptomic and proteomic datasets. The table reports the Reactome term name, NES, and adjusted p-value for each pathway. Pathways are color-coded consistently between the table, enrichment curves, and barcode tracks.

We also examined pathways concordantly depleted in tiR5-high tumors. Among Hallmark gene sets, three pathways were negatively enriched across both transcriptomic and proteomic datasets: protein secretion, MTORC1 signaling, and mitotic spindle (**Fig. S4**). Reactome GSEA identified 71 concordantly negatively enriched pathways shared between transcriptomic and proteomic datasets (**Fig. S5A**). These pathways were most frequently related to cell cycle and DNA replication, which represented the largest category of shared negatively enriched Reactome terms (20 pathways; **Fig. S5B**). Representative depleted pathways included G2/M transition, G1/S transition, synthesis of DNA, DNA replication, and APC/C-mediated mitotic protein degradation (**Fig. S5C**). Other negatively enriched Reactome pathways spanned neuronal, immune, trafficking, morphogenic, metabolic, and protein-turnover programs (**Fig. S5B**).

Overall, these results identify a cross-omics molecular signature of tiR5-high GBM tumors characterized by positive enrichment of EMT, angiogenesis, collagen-related, and ECM remodeling pathways, together with negative enrichment of cell-cycle and DNA-replication programs. This pattern links elevated tiR5 expression to an invasion-associated molecular state that is distinct from canonical proliferative programs. The full GSEA outputs for tiR5-high versus tiR5-low GBM tumor comparisons are provided in **Dataset S3**.

### Elevated tiR5 expression is associated with worse overall patient survival across two independent GBM cohorts

Given the association between elevated tiR5 expression and persistent disease after first-line therapy, we finally asked whether tiR5 expression was associated with worse overall patient survival. Patients were stratified into tiR5-high and tiR5-low groups using mean total tiR5 expression, and survival was evaluated separately in the discovery and confirmatory cohorts. In the discovery cohort (N = 90), tiR5-high tumors were associated with significantly worse overall patient survival compared with tiR5-low tumors (log-rank p = 0.011; **Fig. 8A**). This association was reproduced in the confirmatory cohort (N = 56), where tiR5-high tumors were again associated with worse overall patient survival (log-rank p = 0.0098; **Fig. 8B**).

**Figure 8.**
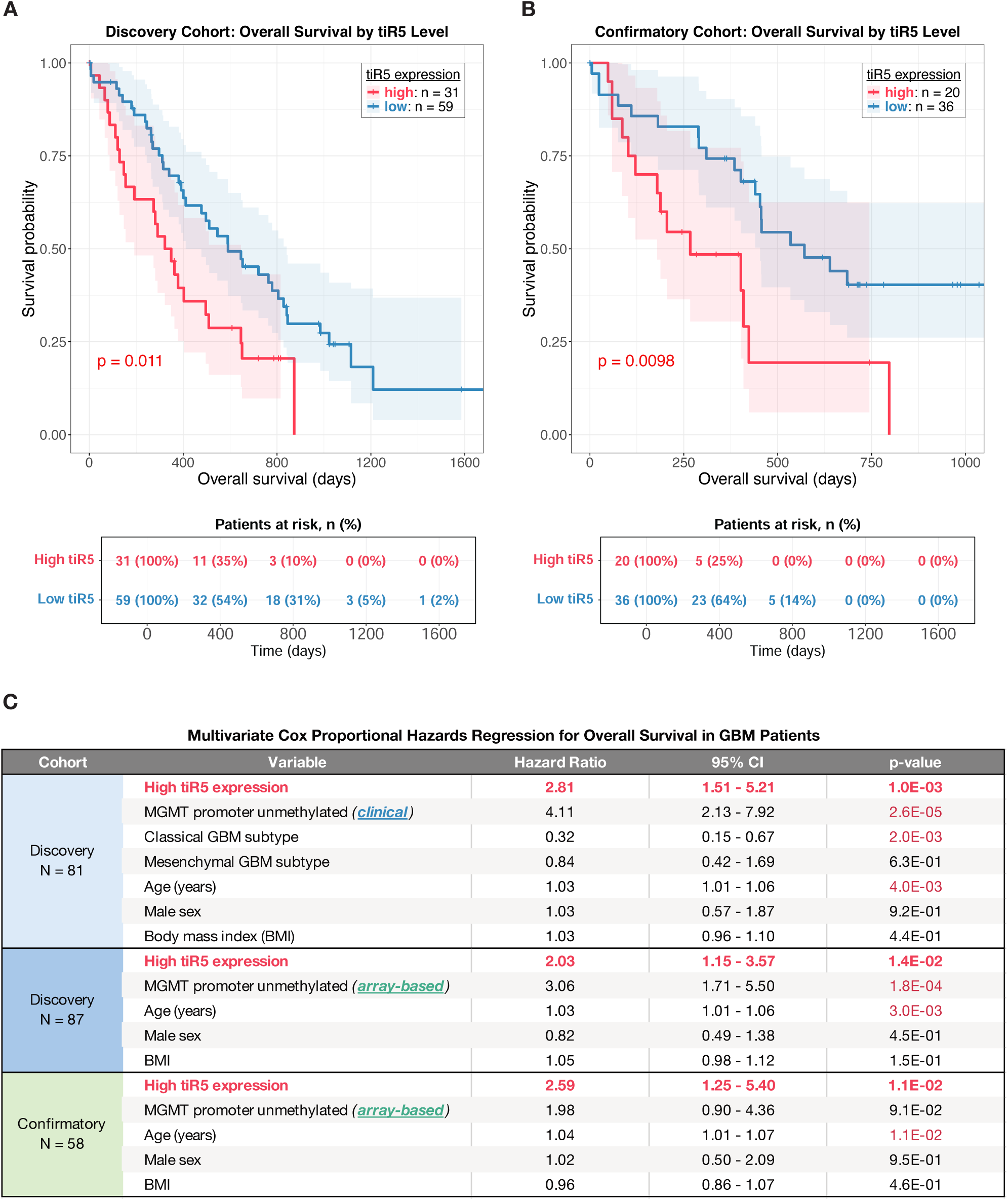
Elevated tiR5 expression is associated with worse overall GBM patient survival across two independent cohorts. (A-B) Kaplan-Meier curves showing overall survival in GBM patients stratified by tiR5 expression level. Patients were classified as tiR5-high or tiR5-low based on mean tiR5 expression. Shaded regions indicate 95% confidence intervals, and tables below each plot show the number and percentage of patients at risk over time. P-values represent log-rank tests comparing tiR5-high and tiR5-low groups. Discovery **(A)** and confirmatory **(B)** cohorts are shown separately. **(C)** Multivariate Cox proportional hazards regression models for overall survival in GBM patients. For the discovery cohort, high tiR5 expression was evaluated with age, sex, BMI, clinical MGMT promoter methylation status, and GBM subtype. To enable cross-cohort comparison, a second model was evaluated in both the discovery and confirmatory cohorts using age, sex, BMI, and array-based MGMT promoter methylation status. Significant p-values ≤ 0.05 are highlighted in red.

We next tested whether high tiR5 expression remained associated with poor patient survival after adjustment for clinical and molecular covariates. In the discovery cohort (N = 81), high tiR5 expression was independently associated with increased mortality risk after adjustment for age, sex, BMI, MGMT promoter methylation status, and GBM subtype (hazard ratio [HR] = 2.81; 95% CI, 1.51-5.21; p = 1.0 x 10^-3^; **Fig. 8C**). MGMT promoter methylation status and age were also significantly associated with survival in this model, whereas classical GBM subtype was associated with lower mortality risk.

We then evaluated whether this survival association was reproducible in the confirmatory cohort. Since clinical MGMT promoter methylation status and GBM subtype were unavailable for the confirmatory cohort, we used a shared covariate model that included age, sex, BMI, and array-based MGMT promoter methylation status. In the discovery cohort (N = 87), high tiR5 expression remained significantly associated with worse overall survival in the shared model (HR = 2.03; 95% CI, 1.15-3.57; p = 1.4 x 10^-2^). In the confirmatory cohort (N = 58), high tiR5 expression was similarly associated with worse overall survival after adjustment for the same covariates (HR = 2.59; 95% CI, 1.25-5.40; p = 1.1 x 10^-2^) (**Fig. 8C**). Across models, patients with tiR5-high tumors had approximately two-to three-fold higher mortality risk than patients with tiR5-low tumors after covariate adjustment.

Together, these results demonstrate that elevated tiR5 expression is associated with worse overall survival across two independent GBM cohorts and remains prognostic after adjustment for established clinical and molecular variables. Combined with the preceding clinical, experimental, and molecular analyses, these findings link elevated tiR5 expression to persistent disease, experimental radiation resistance, pro-invasive molecular programs, and poor survival outcomes in primary patient GBM. These results identify tiR5s as a previously underrecognized small RNA feature associated with aggressive GBM biology and adverse patient outcomes.

## Discussion

In this study, we identify tiR5s as a dominant component of the GBM small RNA landscape and show that elevated tiR5 expression consistently marks aggressive disease across independent patient cohorts and experimental models. Together, the clinical, molecular, and experimental data support a model in which tiR5 expression identifies a stress-adapted GBM state associated with therapeutic resistance, invasive biology, and poor patient outcomes.

Our initial GBM tumor versus non-tumor comparisons revealed coordinated tRF-family level shifts. Specifically, tRF1, tRF3-CCA, tiR5, and tRF leader species were significantly increased in GBM tumors compared to non-tumor brain samples, while tRF5 and tRF3 species were decreased (**Fig. 1C-D**; **Fig. S1**). The coordinated nature of these shifts argues against random tRNA degradation. Rather, this pattern may reflect altered activity, localization, and RNA-modification dependence of tRNA processing enzymes. tRF biogenesis requires site-specific tRNA cleavage by distinct tRNA processing enzymes, a process significantly regulated by tRNA modification status.^55,56^ For example, angiogenin (ANG) is known to cleave tRNAs at the anticodon loop to generate tiRs, yet stress-induced ANG relocalization^34,57–59^ and tRNA modifications^60–62^ often regulate tiR biogenesis independently of changes in total ANG expression. Future studies should therefore evaluate the expression and subcellular localization of tRNA processing enzymes together with tRNA modification patterns in GBM tumors to define the mechanism underlying GBM-associated tRF changes.

Our findings link highly abundant tiR5 species to treatment response within primary GBM, extending two prior glioma tRF profiling studies. Three of the six dominant tiR5s in our analyses, Gly-GCC-tiR5, Gly-CCC-tiR5, and Lys-CTT-tiR5, were previously reported as significantly increased in GBM compared with LGG patient tumors.^48^ A second study also found tiR5s to be highly abundant in GBM and LGG tumors but did not observe significant increases in their three target tiR5 species, Gly-GCC-tiR5, Glu-CTC-tiR5, and Gly-CCC-tiR5, in GBM versus LGG comparisons. However, the same study reported strong upregulation of these three tiR5s across three human GBM cell lines compared with normal astrocyte cells.^47^ While neither study included non-tumor brain comparisons, both studies agreed that the proportion of tRF reads mapped to tiR5s in GBM tumors exceeded that of LGG tumors.^47,48^ These prior findings bear conceptual resemblance to our observations that tiR5s are increased in GBM tumors compared with non-tumor brain (**Fig. 1D**), enriched in GBM tumors that later developed persistent disease (**Fig. 3C-F**), and further increased in radiation-resistant PDX models (**Fig. 5C-D**). The recurrence of these same tiR5 species across independent patient glioma comparisons^47,48^ together with their strong co-expression (**Fig. S2C-D**) and dominance over the tiR5 pool (**Fig. 2D**; **Fig. 5B**; **Fig. S2B**) suggests that dominant tiR5s may mark aggressive glioma biology rather than a single clinical contrast. Together, these findings raise the possibility that tiR5 abundance tracks with increasing glioma aggressiveness across non-tumor brain, LGG, GBM, and treatment-resistant GBM. This possibility remains to be tested directly across matched disease states.

tiR5s remained the dominant signal in our analyses, with high abundance in GBM tumors (**Fig. 2A**; **Fig. 5B**; **Fig. S2A**), increased expression in radiation-resistant GBM PDX models (**Fig. 5C-D**), and associations with persistent disease and worse overall survival across patient cohorts (**Fig. 3C-F**; **Fig. 4**; **Fig. 8**). This association between tiR5s and persistent disease is consistent with prior work linking tiR5s to cancer progression and stress-states. Across cancer types, tiR5s have been shown to regulate tumor growth,^19^ migration,^36^ hypoxia-associated progression,^37^ RNA processing,^36^ and tumor-suppressive signaling.^38^ In our analyses, tiR5-high tumors were enriched for mitochondrial translation programs at the proteomic level (**Fig. 6**), whereas literature supports that pharmacologic inhibition of mitochondrial translation reduces cell growth, clonogenicity, and promotes apoptosis of human GBM stem cells.^63^ Likewise, our analyses showed that tiR5-high tumors displayed concordant transcriptomic and proteomic enrichment of angiogenesis, EMT, and ECM-remodeling programs, with concurrent depletion of cell-cycle and DNA-replication programs (**Fig. 7**; **Fig. S4**; **Fig. S5**). These patterns suggest that elevated tiR5 expression occurs in tumors with pro-invasive and less proliferative molecular profiles. In GBM, such profiles could explain the association between elevated tiR5 expression and poor treatment response. A highly invasive, less proliferative tumor may leave behind diffuse residual disease that extends beyond surgical margins and may be less vulnerable to therapies that preferentially target rapidly dividing tumor cells, including TMZ and radiation. Further studies will be needed to determine whether tiR5s directly contribute to GBM therapeutic resistance or primarily serve as prognostic biomarkers.

Mechanistically, tiR5s are unlikely to act through canonical miRNA-like repression. The tiR5s detected in our analyses were approximately 32-33 nucleotides long, which exceeds the established size range for Argonaute-loaded tRFs (**Fig. 2C**).^16,64^ Instead, tiR5s have been shown to regulate stress-adaptive translational repression, stress granule formation, and anti-apoptotic signaling through mechanisms distinct from miRNA-like targeting, often through interactions with RNA-binding proteins.^12–14^ These stress-adaptive pathways may be relevant to GBM, where tumor cells must withstand hypoxia, nutrient limitation, and treatment-induced toxicity to survive. Further studies manipulating tiR5 species and defining their RNA-binding protein partners will help establish whether tiR5s directly mediate GBM invasiveness, proliferation, and treatment sensitivity in preclinical models.

Beyond potential biological roles, our results support tiR5s as candidate prognostic biomarkers in GBM. Our results showed that tumor tiR5 abundance significantly improves therapeutic response predictions and stratifies patient survival across two independent cohorts (**Fig. 4**; **Fig. 8**). The routine collection and molecular analysis of GBM tissue after initial surgical resection makes tumor-based tiR5 profiling feasible in clinical practice. As further support, tiR5s have been studied as candidate diagnostic and prognostic biomarkers across multiple cancers, with evidence from both tumor tissue and peripheral blood samples.^39–45^ Additionally, tiR5s are readily detected in minimally invasive biofluids, including serum, plasma, and extracellular vesicles.^65–72^ In glioma, tiR5s were previously identified as a prominent feature of glioma stem cell-derived extracellular RNA, including Gly-GCC-tiR5 and Glu-CTC-tiR5, the two most abundant tiR5 species in our GBM cohorts.^49^ These findings support two translational directions: testing tiR5s as tumor-tissue biomarkers in routinely collected GBM specimens and evaluating GBM-associated tiR5s in minimally invasive biofluids for early detection or pre-treatment risk stratification.

Although tiR5s were the focus of this study, the relative abundance of tiR3s should be interpreted cautiously. tiR5s and tiR3s are reciprocal halves of parental tRNAs, but tiR5s are often detected at much higher levels than tiR3s by standard small RNA-sequencing.^26,31,46,48,66,70,73–77^ This apparent imbalance may reflect library-preparation bias, rather than true tiR3 depletion, as 3’-aminoacylation and base-pair disrupting RNA modifications have been shown to impede adapter ligation and reverse transcription of tRNAs and tRFs.^78–83^ tRF detection by northern blot also produces comparable tiR3 and tiR5 signals, suggesting that small RNA-sequencing may exaggerate the difference between the two halves.^19,34,35,62,68^ In our analyses, many tiR3 species were similarly increased in tumors that developed persistent disease following therapy compared with tumors that underwent complete remission (**Fig. 3B-D**). Total tiR3 expression was also strongly correlated with total tiR5 expression in both cohorts (**Fig. S2E-F**), suggesting that tiR3s are produced alongside tiR5s in GBM despite their low apparent abundance. Notably, tiR5 predominance was still observed in our PDX models (**Fig. 5B**), which were profiled using Induro-seq, a library preparation method that aims to minimize RNA modification-associated detection bias.^84^ Therefore, the low apparent tiR3 abundance in GBM tumors may reflect true tiR3 depletion, residual technical bias, or both. These technicalities led us to primarily investigate tiR5s, but our findings do not rule out the potential relevance of tiR3s in GBM treatment resistance. Future studies using sequencing methods fully optimized for modified or aminoacylated tRFs, along with orthogonal northern blot validation will be needed to fully investigate tiR3s in GBM tumors.

Overall, our findings establish tiR5s as an underrecognized molecular feature of primary patient GBM and link elevated tiR5 expression to persistent disease, radiation resistance, pro-invasive molecular programs, and poor patient survival. Across patient cohorts, PDX models, and integrated transcriptomic and proteomic analyses, elevated tiR5 expression consistently distinguished GBM tumors with aggressive disease features. These findings expand the study of GBM heterogeneity to include tRNA fragmentation and support further evaluation of tiR5s as tumor biomarkers and candidate functional contributors to GBM therapeutic resistance.

### Limitations

Multiple limitations should be noted. First, the tumor versus non-tumor comparison included a small number of non-tumor brain samples (N = 8), and the confirmatory cohort did not include matched non-tumor brain samples. Independent validation of GBM versus non-tumor tRF remodeling is therefore still needed. Second, treatment-response analyses were retrospective and based on available clinical annotations. These annotations may not fully capture radiographic and histologic uncertainty, extent of resection, or variations in therapeutic doses and timing. Third, standard small RNA-sequencing methods may under-detect specific tRF families, particularly 3’ tRFs, due to RNA modifications and 3’-terminal chemistry that can interfere with small RNA library preparation.^78–85^ Fourth, the PDX analysis was limited by the small number of matched models (7 pairs) and incomplete technical replication for some samples. Finally, although tiR5 associations with persistent disease and survival were reproduced across cohorts and supported by radiation-resistant PDX models, the study remains correlative. Functional experiments are needed to determine whether tiR5s directly contribute to GBM biology or instead mark aggressive tumor states.

## Methods

### Lead contact

Further information and requests for resources should be directed to the lead contact, Zhangli Su

### Materials availability

No new unique reagents were generated in this study.

### Data and code availability

CPTAC-3 GBM discovery and confirmatory cohort small RNA-sequencing, bulk RNA-sequencing, DNA methylation array, and clinical annotation data were obtained from Genomic Data Commons (GDC).^50^ Discovery cohort proteomic MS1 spectral intensity data were obtained from the CPTAC Pan-Cancer Proteogenomics project in Proteomic Data Commons (PDC), using the harmonized Baylor College of Medicine proteome dataset titled Proteome_BCM_GENCODE_v34_harmonized_v1.zip.^54^ Additional CPTAC-3 GBM discovery cohort clinical and molecular annotations were obtained using the CPTAC Python package.^86^ PDX small RNA-sequencing data generated for this study will be deposited in the Gene Expression Omnibus (GEO) under accession GSE338531.

### Experimental model and study participant details

#### CPTAC-3 patient cohorts and samples

Publicly available multi-omic datasets from two CPTAC-3 GBM cohorts, discovery and confirmatory, were used for this study. The discovery cohort included 92 resected primary GBM tumor samples, each from an individual patient, collected before chemoradiotherapy. The discovery cohort also included 8 age-matched post-mortem non-tumor brain samples from GTEx that were processed alongside the GBM samples by the CPTAC group.^50^ The confirmatory cohort included 67 primary GBM tumor samples, each from an individual patient, and did not include non-tumor brain samples. IDH-mutant glioma samples were excluded from all analyses. General cohort characteristics are described in **Table 1**.

#### GBM PDX models

Total RNA from GBM PDX models was obtained from the University of Alabama at Birmingham (UAB). The initial radiation-sensitivity screening and *in vivo* selection of radiation-resistant GBM PDX models were described previously.^53^ The original patient-derived GBM tissues were obtained from the UAB Brain Tumor Tissue Core and from JN Sarkaria at Mayo Clinic (Rochester, MN, USA). The final small RNA-seq dataset included 7 matched baseline and radiation-resistant PDX pairs, with up to 3 technical replicates per condition. RNA integrity was assessed using a Bioanalyzer, and samples showed acceptable RNA quality overall, with a median RIN of 7.15 (SD = 1.62). Although 3 technical replicates were planned for each PDX line and condition, the study used residual RNA from previously generated PDX samples, and sufficient RNA was not available for all planned replicates. The final dataset included 19 baseline and 16 radiation-resistant RNA samples.

## Method details

### Generation of GBM PDX models

GBM PDX models were generated as described previously.^53^ Briefly, patient-derived GBM xenografts were established as flank tumors in athymic nude mice. Tumors were irradiated when they reached 200-300 mm^3^. Local tumor irradiation was delivered as 2 Gy per fraction on Monday, Wednesday, and Friday for 2 weeks, for a total dose of 12 Gy. Radiation response was defined by time to tumor-size doubling after the first radiation dose. Tumors were classified as radiation-resistant if tumor size doubled within 20 days of initial irradiation (**Fig. 5A**).

### Small RNA-seq library preparation from GBM PDX models

Total RNA was isolated from frozen GBM PDX tissue using TRIzol reagent followed by chloroform phase separation, isopropanol precipitation, 75% ethanol wash, and resuspension in nuclease-free water. Small RNA-seq libraries were prepared from total RNA using Induro-seq, a modified protocol utilizing the commercially available NEBNext Small RNA Library Prep Set for Illumina (New England Biolabs, Cat. #E7330), replacing the reverse transcriptase with Induro-RT (New England Biolabs, Cat. #M0681S).^84^ Unlike the referenced protocol, subcellular fractionation was omitted and libraries were prepared directly using 100 ng of total RNA. Indexing was performed using NEBNext Multiplex Oligos for Illumina (New England Biolabs, Cat. #E6440S). Following adapter ligation, indexing, reverse transcription, and amplification, library products were size-selected by excision from an 8% polyacrylamide gel (Thermo Fisher Scientific, Cat. #EC6215BOX) to enrich for inserts derived from RNAs ≤ 50 nt, pooled, and sequenced on Illumina NovaSeq X Plus (Novogene).

### Quantification and statistical analysis

All statistical analyses and visualizations were performed using R (v4.4.1) in RStudio (release 2026.05). All included p-values were adjusted using the Benjamini-Hochberg method unless otherwise specified.

### Clinical and molecular annotations

Clinical and molecular annotations were obtained from GDC data portals and the CPTAC-3 Python package.^86^ Treatment response was assigned directly from the CPTAC-3 clinical annotation field “measure_of_success_of_outcome_at_the_completion_of_initial_first_course_treatment,” where tumors were annotated as either “complete remission” or “persistent disease.” Overall survival was analyzed in days, and survival status was coded as 1 for deceased and 0 for alive. For discovery cohort tumors, stromal score, immune score, and tumor purity fields were obtained from ESTIMATE^87^ annotations provided by CPTAC.

### Small RNA-seq processing and tRF annotation

For CPTAC-3 patient samples, GDC small RNA-seq BAM files were converted to FASTQ format using bedtools (v2.31.1).^88^ Adapter trimming had been performed by CPTAC before file download. For PDX samples, raw FASTQ files generated from Induro-seq libraries were adapter-trimmed using cutadapt (v5.2).^89^ Trimmed small-RNA reads from CPTAC-3 patient and PDX datasets were mapped and annotated using unitas (v1.7.3), a small RNA annotation tool that classifies reads across multiple RNA biotypes, including tRFs.^51^ Unitas performs sequential mapping, first to human miRBase (release 22),^90^ followed by other small RNA reference databases, including GtRNAdb (release 2.0),^91^ SILVA rRNA database (release 132),^92^ and Ensembl *Homo sapiens* annotations (release 97). Multi-mapped reads were fractionally assigned by dividing each read count equally across all compatible annotations.

### Count normalization and tiR5 quantification

Small-RNA features with a summed raw count of fewer than 10 reads across all samples were removed before downstream analysis. Remaining counts were normalized using DESeq2 (v1.44.0).^93^ Unless otherwise specified, DESeq2-normalized counts were log_2_-transformed after adding a pseudocount of 1.

Total tiR5 expression was calculated for each sample by summing DESeq2-normalized counts across all annotated tiR5 species. Summed tiR5 counts were then log_2_-transformed after adding a pseudocount of 1 unless otherwise noted. For tiR5-high and tiR5-low analyses, samples were classified using the cohort-specific mean total tiR5 expression. Samples with total tiR5 expression greater than or equal to the cohort mean were classified as tiR5-high, and samples below the cohort mean were classified as tiR5-low.

Proportional abundance analyses were calculated from mapped read counts after DESeq2 normalization. RNA biotype, tRF family, and individual tiR5 species proportions were calculated as percentages of total mapped small-RNA reads, total mapped tRF reads, and total tiR5 reads, respectively (**Fig. 1A; Fig. 2A** and **D**; **Fig. S2A-B**). tiR5 read-length distributions were calculated from unitas-annotated tiR5 reads before DESeq2 normalization (**Fig. 2C**). Visualizations were generated using ggplot2 (v4.0.2) and ggpubr (v0.6.0).

### Small RNA differential expression analyses

Differential expression between GBM tumors and non-tumor brain samples was performed in the discovery cohort using DESeq2 (v1.44.0),^93^ comparing tumors (N = 92) with non-tumor samples (N = 8) using a design formula containing sample group only (**Fig. 1C**; **Fig. S1**). For tumor versus non-tumor family-level fold-change distribution analyses, log_2_ fold-change values for all detected members of each RNA class or tRF family were compared with the log_2_ fold-change distribution across all small RNAs shown in the panel (**Fig. 1D**). Comparisons were performed using Wilcoxon rank-sum tests followed by p-value correction.

Clinical and molecular feature screening was performed in the discovery cohort using all samples with available annotations for each comparison. Binary features were analyzed using DESeq2 (v1.44.0),^93^ whereas features with more than two groups were analyzed using Kruskal-Wallis tests. Screened features included GBM subtype, NMF subtype, tumor recurrence, telomere length, MGMT promoter methylation, chromosome 7 amplification, chromosome 10 deletion, and treatment response (**Fig. 3B**; **Fig. S3**). Treatment-response associations were then evaluated in both the discovery and confirmatory cohorts.

For treatment-response differential expression analyses, tumors annotated as persistent disease were compared with tumors annotated as complete remission using DESeq2 (v1.44.0)^93^ with a design formula containing treatment-response group only (**Fig. 3C-D**). Discovery and confirmatory cohorts were analyzed separately. Differentially expressed small RNAs were defined by adjusted p-value ≤ 0.05 and absolute log_2_ fold change ≥ 0.5.

For PDX analyses, resistant-to-baseline fold changes were calculated for total tiR5 expression and for the six dominant tiR5 species within matched PDX model pairs. Fold-change distributions were tested against a null fold change of 1 using Wilcoxon signed-rank tests, followed by p-value correction (**Fig. 5C-D**). Data visualizations were generated using ggplot2 (v4.0.2) and ggpubr (v0.6.0) R packages.

### Correlation and heatmap analyses

Pairwise correlations among the six dominant tiR5 species (**Fig. S2C-D**) and among tRF families (**Fig. S2E-F**) were assessed using Pearson correlation coefficients. P-values were adjusted for multiple testing. Correlations between total tiR5 expression and tissue composition scores were also assessed using Pearson correlation coefficients (**Fig. S2G**). P-values were reported without multiple-testing adjustment for tissue composition analyses since only three comparisons were performed. Correlation heatmaps and scatterplots were generated in R (v4.4.1) using corrplot (v0.94) and ggplot2 (v4.0.2).

Heatmaps were generated from either log_2_-normalized tiR5 expression values or MS1 protein abundance values. Before visualization, values were scaled by row using z-score transformation. For tiR5 heatmaps, rows corresponded to total tiR5 expression or individual tiR5 species (**Fig. S3A**). For proteomic heatmaps, rows corresponded to leading-edge proteins from enriched mitochondrial translation Reactome pathways (**Fig. 6B**). Heatmaps were generated using the pheatmap (v1.0.12) package in R.

### Array-derived MGMT promoter methylation annotation

MGMT promoter methylation was inferred from DNA methylation array data using the two-CpG STP27 approach.^94,95^ Probe annotations were obtained from the EPIC_hg38_annotations.tsv file provided through the GDC community tools webpage. Beta values were extracted for two CpG probes within the MGMT promoter, cg12434587 and cg12981137, which were used for the STP27 classifier. For each probe, beta values were converted to M-values using the formula log_2_(beta / [1 - beta]). Before transformation, beta values were bounded to avoid undefined or infinite M-values. The mean M-value across the two MGMT promoter probes was then calculated for each sample.

A cohort-specific methylation cutoff was derived in the discovery cohort using available clinical MGMT promoter methylation annotations as the reference standard. An ROC curve was generated using the mean two-probe M-value as the predictor and clinical MGMT promoter methylation status as the reference label. The cutoff that maximized the Youden index was selected and then applied uniformly to both the discovery and confirmatory cohorts. Samples with mean two-probe M-values greater than or equal to the cutoff were classified as MGMT promoter methylated, whereas samples below the cutoff were classified as unmethylated. Cutoff selection and classifier performance assessment were performed using the pROC (v1.18.5) package in R.

In the discovery cohort, the two-probe array-derived classifier showed high concordance with clinical MGMT promoter methylation annotations, with an AUC of 0.97. Among 83 discovery cohort samples with available clinical MGMT annotations, only 1 sample was assigned a different methylation status by the array-derived classifier.

### ROC analysis for persistent disease

Logistic regression was used to evaluate prediction of persistent disease after first-line therapy. Treatment response was modeled as a binary outcome comparing persistent disease with complete remission. Total tiR5 expression was included as a continuous log_2_-transformed variable. Age and BMI were modeled as continuous variables. Sex was modeled as a categorical variable with female as the reference level. MGMT promoter methylation status was modeled as a categorical variable with methylated as the reference level.

Two clinical models were evaluated. Clinical model A included age, sex, BMI, and clinical MGMT promoter methylation status and was restricted to the discovery cohort, where clinical MGMT annotations were available. Clinical model B included age, sex, BMI, and array-derived MGMT promoter methylation status and was evaluated in both the discovery and confirmatory cohorts (**Fig. 4A**). For each clinical model, three prediction models were tested: clinical variables alone, clinical variables plus tiR5 expression, and tiR5 expression alone.

Model performance was assessed using 5-fold cross-validation repeated 100 times. Within each repeat, logistic regression models were fit in the training folds and used to generate predicted probabilities for held-out samples. Out-of-fold predictions were averaged across repeats for each patient. ROC curves and AUC values with 95% confidence intervals were then calculated for each model. AUCs for each tiR5-containing model were compared against base clinical models using one-sided DeLong’s tests, with the alternative hypothesis being that adding tiR5 expression increased AUC (**Fig. 4B-E**). All ROC analyses and visualizations were performed using the caret (v7.0-1) and pROC (v1.18.5) packages in R.

### Transcriptomic and proteomic GSEA

Discovery cohort bulk RNA-seq data were obtained from GDC as STAR-aligned transcript-level count files, and discovery cohort proteomic data were obtained from PDC as a harmonized log_2_-transformed MS1 spectral intensity matrix. Patient tumors (N = 92) were stratified by total tiR5 expression using the mean total tiR5 expression across the discovery cohort. Samples with total tiR5 expression greater than or equal to this mean were classified as tiR5-high, whereas samples below this mean were classified as tiR5-low.

For the transcriptomic analysis, transcripts with fewer than 10 total raw counts across all samples were removed before downstream analysis. Differential expression analysis was performed using DESeq2 (v1.44.0),^93^ comparing tiR5-high tumors with tiR5-low tumors. For the proteomic analysis, protein abundance was compared between tiR5-high and tiR5-low tumors using Wilcoxon rank-sum tests (**Fig. 6**).

For each data type, the resulting feature-level statistics were used to generate a ranked list for GSEA. Ranking statistics were calculated as-log_10_(p-value) x sign(log_2_ fold change). Positive ranking values indicated higher RNA expression or protein abundance in tiR5-high tumors, whereas negative ranking values indicated higher RNA expression or protein abundance in tiR5-low tumors. GSEA was performed separately for the transcriptomic and proteomic ranked lists using Reactome and MSigDB Hallmark gene sets.

Concordant transcriptomic and proteomic pathways were identified by overlapping GSEA terms that were significant in both data types. Significance was defined as an adjusted p-value ≤ 0.05. Concordant pathways were further classified by enrichment direction, with positive concordance defined by positive NES values in both analyses (**Fig. 7**) and negative concordance defined by negative NES values in both analyses (**Fig. S4**; **Fig. S5**).

GSEA was performed using the clusterProfiler (v4.12.6), msigdbr (v25.1.1), and ReactomePA (v1.48.0) packages in R. Gene symbols were converted to Entrez identifiers using the org.Hs.eg.db (v3.19.1) package. GSEA visualizations were generated using the enrichplot (v1.24.4) package, and pathway overlaps were visualized using the ggvenn (v0.1.10) package.

### Survival analysis

Overall survival was evaluated in the discovery and confirmatory CPTAC-3 GBM cohorts. Within each cohort, patients were stratified by total tiR5 expression using the cohort-specific mean. Samples with total tiR5 expression greater than or equal to the cohort mean were classified as tiR5-high, whereas samples below the cohort mean were classified as tiR5-low. Kaplan-Meier survival analysis was performed to compare overall survival between tiR5-high and tiR5-low groups in the discovery cohort (N = 90) and confirmatory cohort (N = 56). Survival curves were compared using log-rank tests and were plotted separately for each cohort (**Fig. 8A-B**).

Multivariable Cox proportional hazards regression was used to test whether high tiR5 expression was associated with overall survival after adjustment for clinical and molecular covariates (**Fig. 8C**). In the discovery cohort, the first model included tiR5 level, clinical MGMT promoter methylation status, GBM subtype, age, sex, and BMI (N = 81). To enable comparison across cohorts using shared covariates, an additional model was fit in the discovery cohort (N = 87) and confirmatory cohort (N = 58) using tiR5 level, array-derived MGMT promoter methylation status, age, sex, and BMI.

In Cox models, tiR5 level, MGMT promoter methylation status, GBM subtype, and sex were modeled as categorical variables. Reference levels were tiR5-low, MGMT methylated, proneural subtype, and female sex, respectively. Age and BMI were modeled as continuous variables. Hazard ratios, 95% confidence intervals, and Wald p-values were calculated for each covariate (**Fig. 8C**). Survival analyses were performed in R using survival (v3.7-0) and survminer (v0.4.9).

## Acknowledgements

This work was supported by the UAB O’Neal Invests Program Catalyst Award (to Z.S., C.D.W., and E.Y.A.), NIH grants R00 CA259526 (to Z.S.), T32 NS121721 (to M.A. and T.S.), and T32 GM135028 (to S.D.). Generation of radiation-resistant PDX models was supported by NIH grants U01 CA223976 and U01 CA223976-03S1. The authors thank the UAB IT Research Computing team for next-generation sequencing support. BioRender was used to prepare the graphical abstract, **Fig. 3A**, and **Fig. 5A**.

## Author contributions

M.A. performed all bioinformatic analyses, interpreted the results, generated the figures, and wrote the manuscript. Z.S. conceived and supervised the study, provided bioinformatic guidance, and prepared the Induro-seq libraries. T.S. generated the GBM PDX models and collected tumor tissues under the supervision of C.D.W., who developed the PDX irradiation protocol. S.D. and M.M. isolated RNA from frozen PDX tumors under the supervision of E.Y.A. C.D.W., E.Y.A., and Z.S. reviewed and edited the manuscript. All authors reviewed and approved the final manuscript.

---

**Figure S1.**
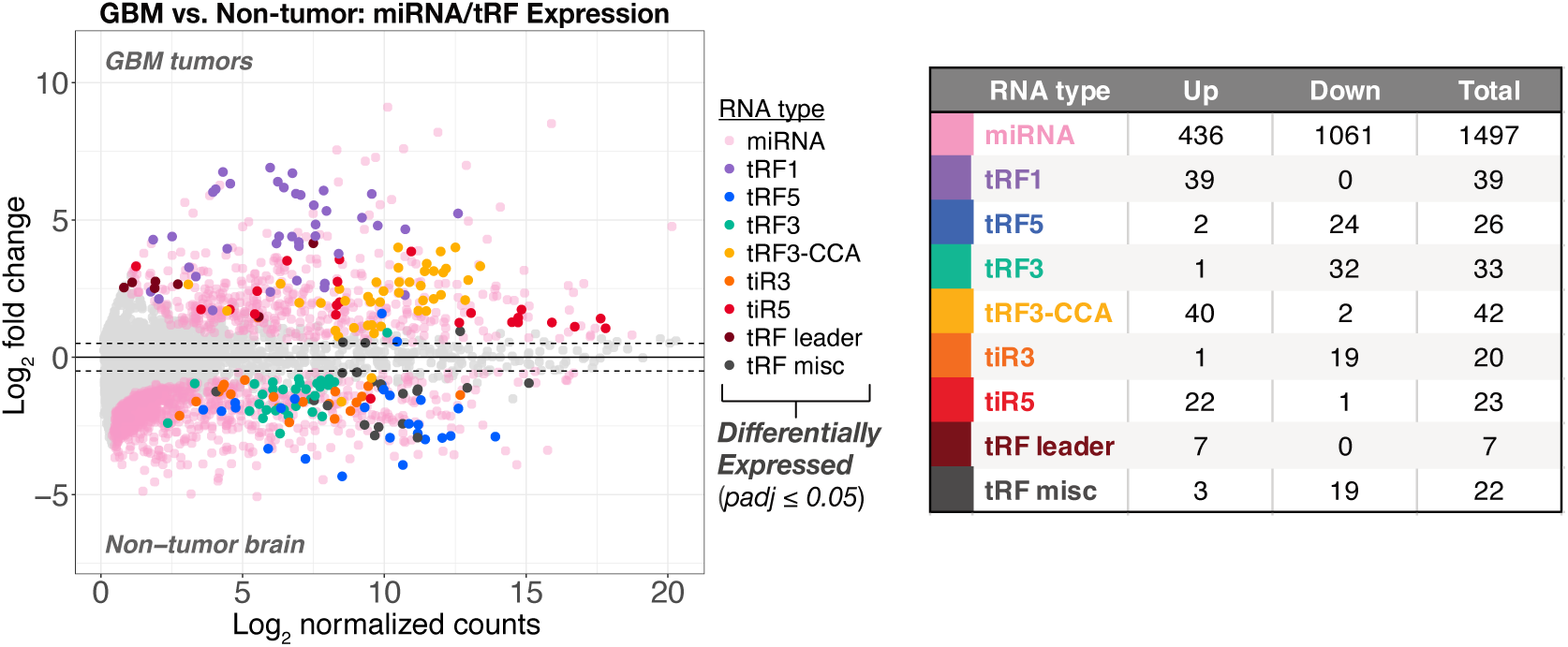
Additional differentially expressed small RNAs in GBM tumors compared with non-tumor brain. MA plot showing differential expression of miRNAs and tRFs in GBM tumors (n = 92) compared with non-tumor brain samples (n = 8). The x-axis shows mean DESeq2-normalized expression on a log_2_ scale, and the y-axis shows log_2_ fold change in GBM tumors versus non-tumor brain. Significantly differentially expressed miRNAs and tRFs are highlighted using an adjusted p-value ≤ 0.05 and an absolute log_2_ fold change ≥ 0.5. Highlighted points are color-coded by miRNA or tRF family. Gray points indicate small RNAs that did not meet these thresholds. The accompanying table reports the number of significantly upregulated and downregulated small RNAs in each RNA class.

**Figure S2.**
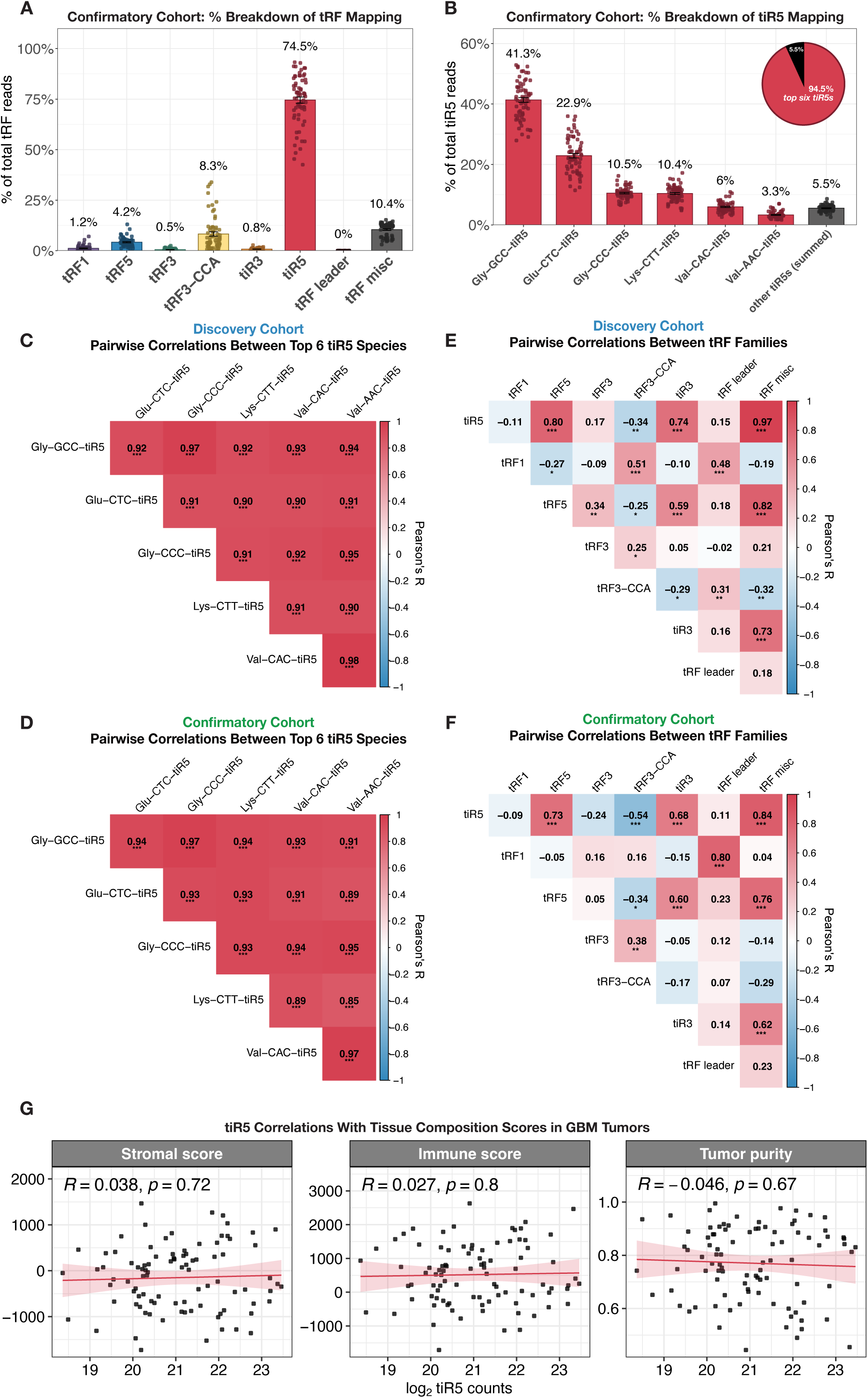
tiR5 abundance, co-expression, and tumor composition analyses in GBM tumors. **(A)** Bar plot showing the percentage of mapped tRF reads assigned to each tRF family in primary GBM tumors from the confirmatory patient cohort (N = 67). **(B)** Bar plot showing the percentage of total tiR5 reads assigned to the six most abundant tiR5 species, with all remaining tiR5 species grouped as “other tiR5s.” The inset pie chart shows that the six most abundant tiR5 species account for 94.5% of total tiR5 reads. For panels A and B, bars show the mean, error bars show SEM, and points represent individual tumors. **(C-D)** Correlation matrices showing pairwise co-expression among the six most abundant tiR5 species in the discovery **(C)** and confirmatory **(D)** cohorts**. (E-F)** Correlation matrices showing pairwise relationships between total tRF family expression levels in the discovery **(E)** and confirmatory **(F)** cohorts. Each cell in panels C-F includes the Pearson correlation coefficient and asterisks that indicate adjusted p-value levels: *p-adj ≤ 0.05, **p-adj ≤ 0.01, and ***p-adj ≤ 0.001. **(G)** Scatter plots showing associations between total tiR5 expression and tumor composition scores in GBM tumors. The x-axis shows log_2_-transformed DESeq2-normalized total tiR5 counts, and the y-axis shows stromal score, immune score, or tumor purity. Pearson’s correlation coefficient and p-value are shown in each plot.

**Figure S3.**
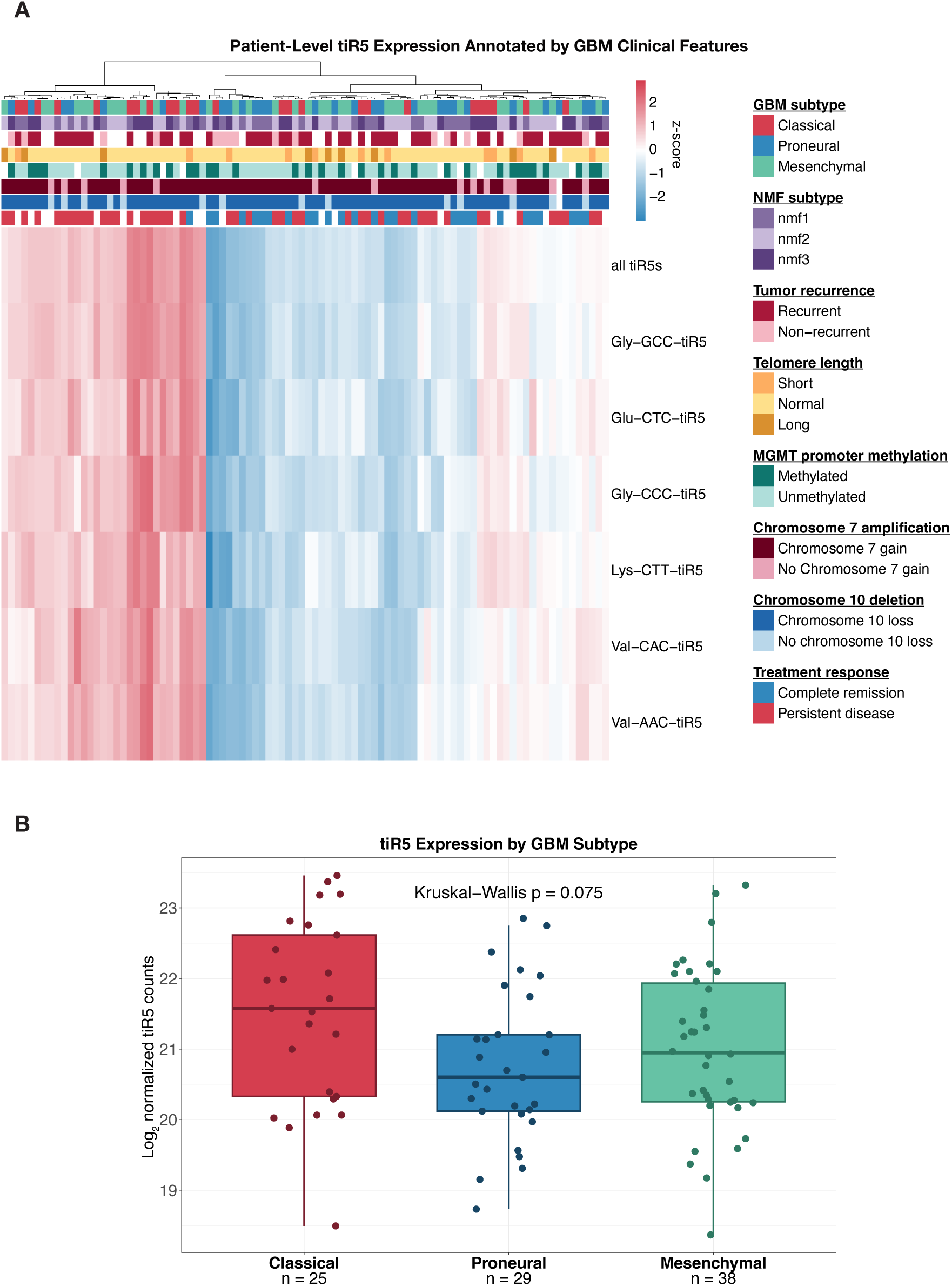
Clinical associations of individual highly expressed and total tiR5s. **(A)** Heatmap showing patient-level expression of total tiR5s and the six most abundant individual tiR5 species in GBM tumors from the discovery cohort. Columns represent individual patient-tumors, and rows represent total or individual tiR5 expression. Samples are clustered by tiR5 expression and annotated by GBM subtype, NMF subtype, tumor recurrence, telomere length, MGMT promoter methylation, chromosome 7 amplification, chromosome 10 deletion, and treatment response. tiR5 expression is shown as a row-scaled z-score across samples. White-colored annotation cells refer to missing phenotype data. **(B)** Boxplot of log_2_-normalized total tiR5 counts across GBM subtypes in the discovery cohort. Each point represents one patient tumor. P-value was calculated using a Kruskal-Wallis test across GBM subtypes.

**Figure S4.**
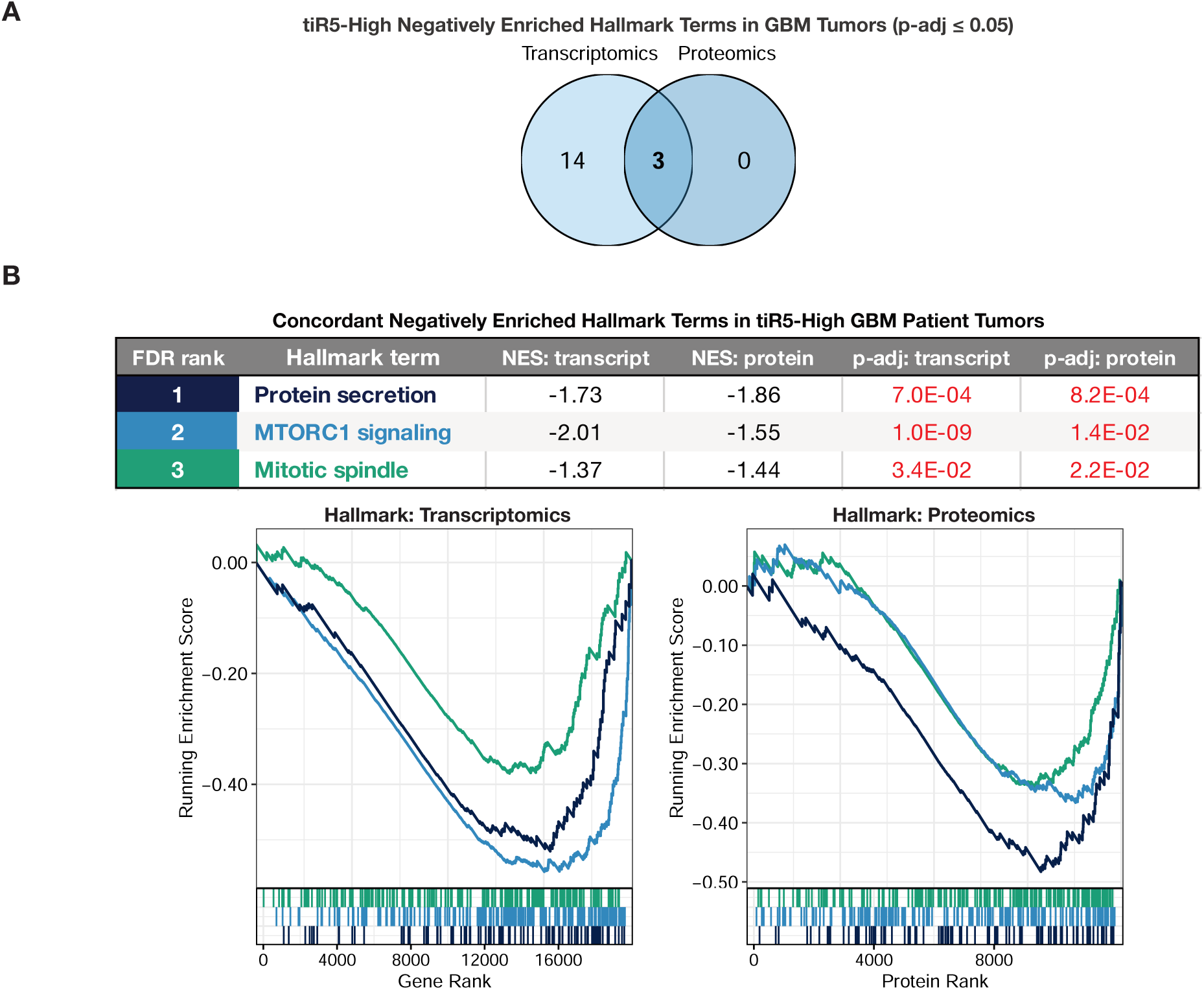
tiR5-high GBM tumors show concordant negative enrichment of protein secretion, mTORC1 signaling, and mitotic spindle pathways. **(A)** Venn diagram showing overlap of significantly negatively enriched MSigDB Hallmark pathways between transcriptomic and proteomic GSEA analyses in tiR5-high GBM tumors. Significantly enriched pathways were defined using an adjusted p-value ≤ 0.05. **(B)** Table and enrichment curves for concordant negatively enriched Hallmark pathways in tiR5-high GBM tumors across transcriptomic and proteomic datasets. The table reports Hallmark term name, normalized enrichment score (NES), and adjusted p-value for each pathway. Negative NES values indicate lower pathway enrichment in tiR5-high tumors. Pathways are color-coded consistently between the table, enrichment curves, and barcode tracks.

**Figure S5.**
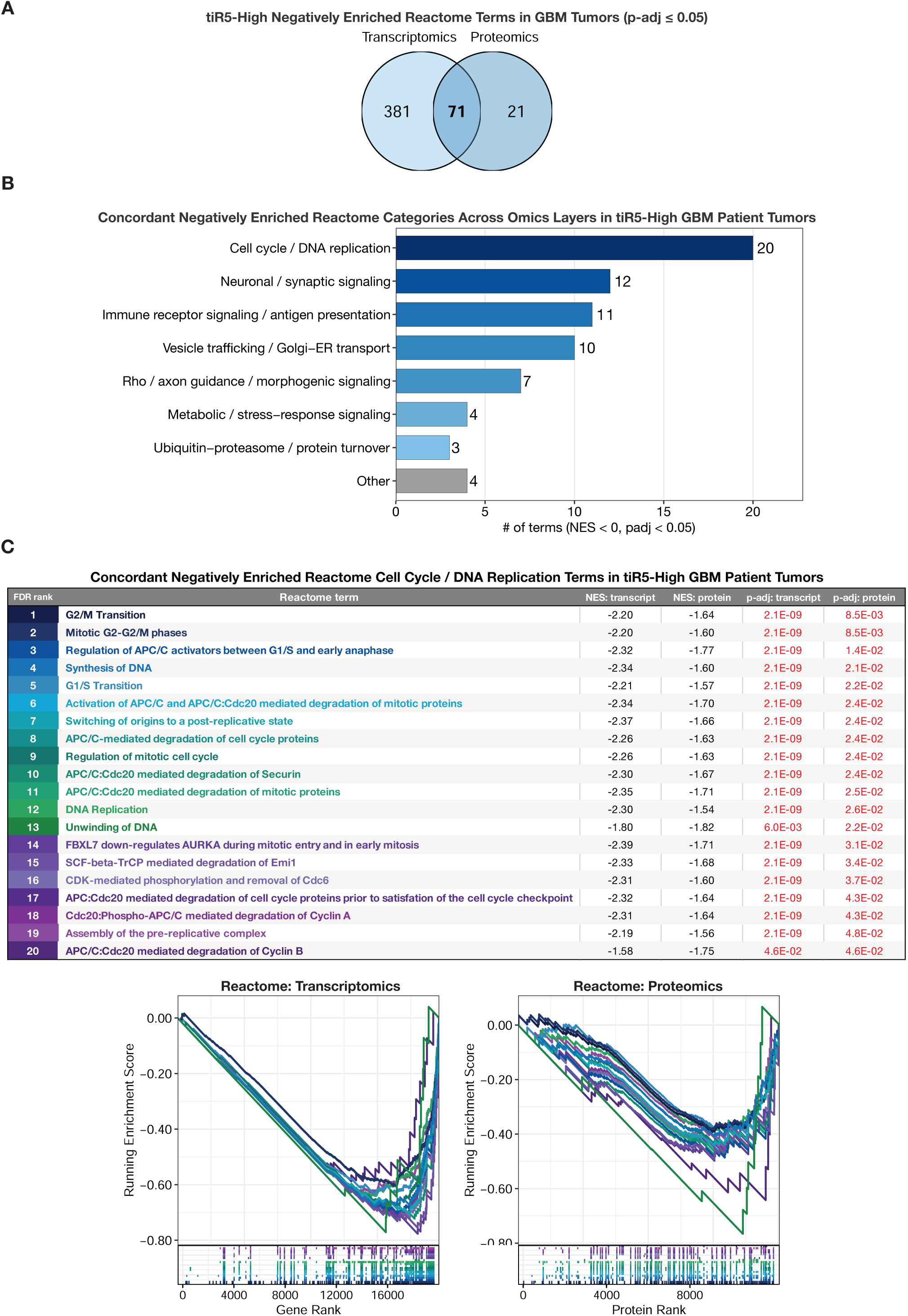
tiR5-high GBM tumors show concordant negative enrichment of cell cycle and DNA replication-related pathways. **(A)** Venn diagram showing overlap of significantly negatively enriched Reactome pathways between transcriptomic and proteomic GSEA analyses in tiR5-high GBM tumors. Significantly enriched pathways were defined using an adjusted p-value ≤ 0.05. **(B)** Bar plot summarizing categories represented among the 71 concordant negatively enriched Reactome pathways shared between transcriptomic and proteomic analyses. The x-axis shows the number of pathways in each category. **(C)** Table and enrichment curves for concordant negatively enriched Reactome cell cycle and DNA replication-related pathways in tiR5-high GBM tumors. The table reports Reactome term name, normalized enrichment score (NES), and adjusted p-value for each pathway. Negative NES values indicate lower pathway enrichment in tiR5-high tumors. Pathways are color-coded consistently between the table, enrichment curves, and barcode tracks.

## Notes

### Competing Interest Statement

The authors have declared no competing interest.

